# The hexosamine biosynthesis pathway is a targetable liability in lung cancers with concurrent KRAS and LKB1 mutations

**DOI:** 10.1101/2020.04.13.039669

**Authors:** Jiyeon Kim, Feng Cai, Bookyung Ko, Chendong Yang, Hyun Min Lee, Nefertiti Muhammad, Kailong Li, Mohamed Haloul, Wen Gu, Brandon Faubert, Akash K. Kaushik, Ling Cai, Sahba Kasiri, Ummay Marriam, Kien Nham, Luc Girard, Hui Wang, Xiankai Sun, James Kim, John D. Minna, Keziban Unsal-Kacmaz, Ralph J. DeBerardinis

## Abstract

In non–small cell lung cancer (NSCLC), concurrent mutations in the oncogene *KRAS* and the tumor suppressor *STK11* encoding the kinase LKB1 result in aggressive tumors prone to metastasis but with liabilities arising from reprogrammed metabolism. We previously demonstrated perturbed nitrogen metabolism and addiction to an unconventional pathway of pyrimidine synthesis in KRAS/LKB1 co-mutant (KL) cancer cells. To gain broader insight into metabolic reprogramming in NSCLC, we analyzed tumor metabolomes in a series of genetically engineered mouse models with oncogenic KRAS combined with mutations in LKB1 or p53. Metabolomics and gene expression profiling pointed towards an activation of the hexosamine biosynthesis pathway (HBP), another nitrogen-related metabolic pathway, in both mouse and human KL mutant tumors. KL cells contain high levels of HBP metabolites, higher flux through the HBP pathway and elevated dependence on the HBP enzyme Glutamine-Fructose-6-Phosphate Transaminase 2 (GFPT2). GFPT2 inhibition selectively reduced KL cell growth in culture and xenografts. Our results define a new metabolic vulnerability in KL tumors and provide a rationale for targeting GFPT2 in this aggressive NSCLC subtype.

## Introduction

Despite recent advances in treatment of non-small cell lung cancers (NSCLCs), they continue to be the leading cause of cancer-related deaths worldwide. Large-scale sequencing efforts have defined the tumor suppressors and oncogenes mutated in NSCLC, but most of these mutations are not yet subject to targeted therapy. Of these mutations, *STK11*, which encodes the tumor suppressor LKB1, is mutated in ∼20% of NSCLCs. *KRAS* is the most frequently mutated oncogene in lung adenocarcinoma (LUAD, ∼30%), the most common histological subtype of NSCLC. Some 6-12% of LUAD contain concurrent mutations in KRAS and LKB1, and these co-mutant tumors display a highly metastatic phenotype not observed with other combinations of mutations^1-3^. LKB1 alterations in the context of KRAS mutation drive primary resistance to PD-1 blockade, rendering these tumors unfavorable for immunotherapy^4^. Thus, further insight into the biology and vulnerabilities of these tumors could yield substantial therapeutic impact.

Metabolic reprogramming is a hallmark of cancer, as tumors alter metabolic pathways to meet the bioenergetic, biosynthetic, and redox requirements of malignancy. Mutation of oncogenes and/or tumor suppressors are a major factor in cell-autonomous metabolic reprogramming. Interestingly, oncogenic KRAS and LKB1 loss independently alter metabolism in cancer cells. Oncogenic KRAS enhances glucose uptake and glycolysis^5,6^, stimulates glutamine catabolism^7^, increases fatty acid uptake for macromolecular synthesis and modulates redox balance^8^. LKB1 participates in fuel sensing by regulating AMP-activated protein kinase (AMPK) and other kinases during energetic stress^9^. The LKB1-AMPK pathway suppresses mTOR complex 1 signaling, re-setting the balance between energy production and utilization^10^. Liabilities in the KRAS/LKB1 co-mutant state include enhanced dependence on the electron transport chain^11^, the pyrimidine synthesis enzyme deoxythymidylate kinase^12^, lysosomal acidification^13^ and the urea cycle enzyme carbamoyl phosphate synthetase 1 (CPS1)^14^.

In order to identify additional metabolic vulnerabilities in KRAS/LKB1 co-mutants, we integrated metabolic and gene expression profiles of lung tumors from a series of genetically-modified mice. This analysis pointed to induction of the hexosamine biosynthesis pathway (HBP) and enhanced reliance on the HBP enzyme Glutamine-Fructose-6-Phosphate Transaminase 2 (GFPT2) in KRAS/LKB1 co-mutant tumors.

## Results

### Lkb1 loss alters metabolism of Kras-mutant lung tumors in mice

We first compared metabolomes between Kras (K), Kras/p53 (KP), and Kras/Lkb1 (KL)-comutant lung tumors and adjacent lung, using autochthonous mouse models from Kwok-Kin Wong’s group^1^ (Supplementary Table 1). Supervised analysis produced the largest number of metabolites differing between KL co-mutant lung tumors and normal lung tissue (Fig 1a). Further, most metabolites that discriminate between K and KL tumors also discriminated between KP and KL tumors, whereas metabolites discriminating between K and KP tumors were largely distinct from ones between K and KL. These data imply that LKB1 mutation imparts a specific metabolomic signature in lung tumors (Supplementary Data 1a,b).

**Figure 1.**
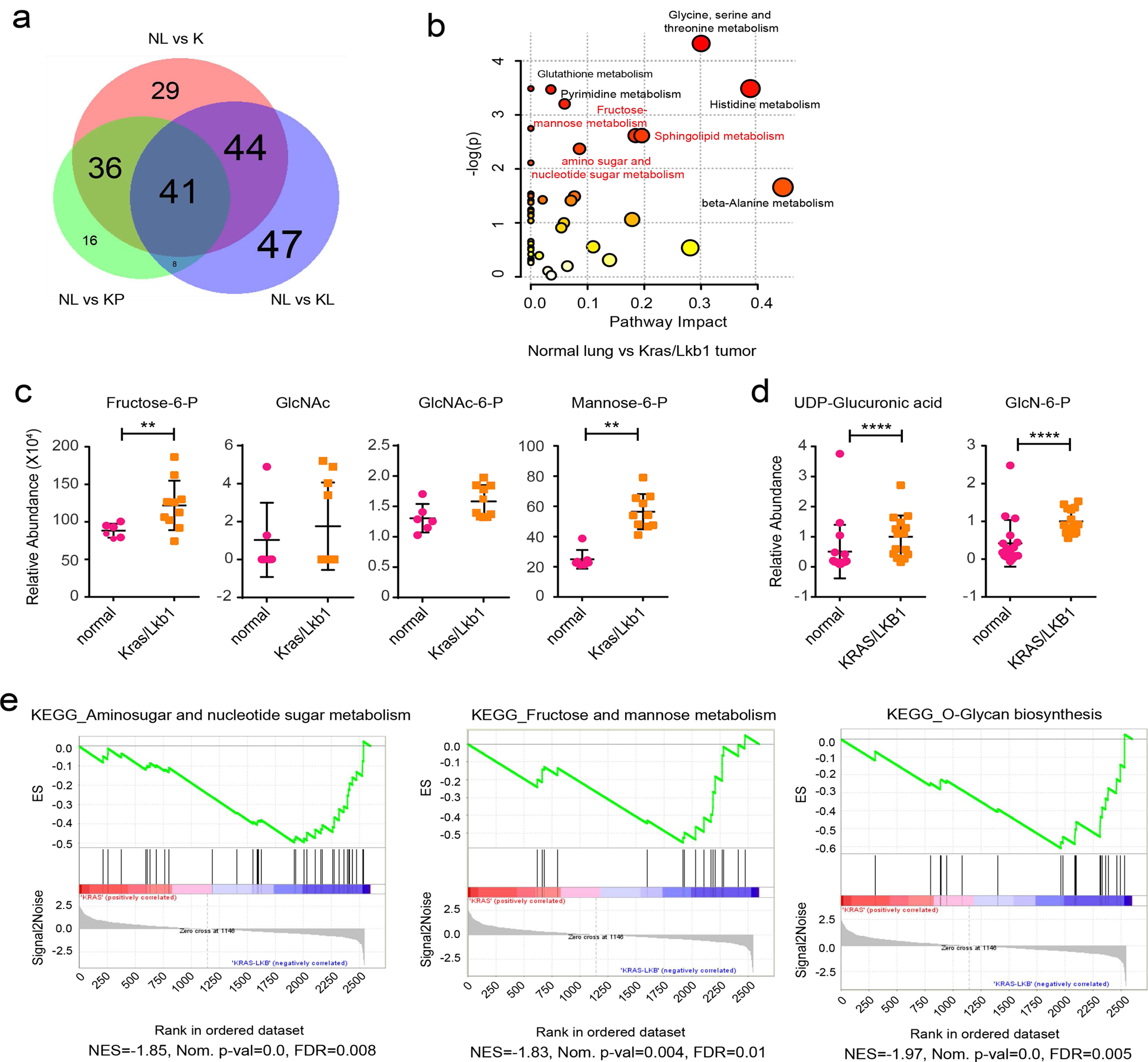
Altered hexosamine metabolism in KL mouse tumors. **a,** Venn diagrams of Variable Importance in the Projection metabolites (VIP>1.0) between normal lung (NL) and tumor tissues with different genotypes. K, Kras mutant tumor; KP, Kras/p53 mutant tumor; KL, Kras/Lkb1 tumor. **b,** Pathway analysis based on metabolites differentiating between normal lung tissues and KL tumor tissues (VIP>1.0). Darkness and size of the dot represents statistical significance of the pathway. Metabolic pathways first observed in the current metabolomics data as opposed to our previous analysis of human tumors^14^ are in red. **c,** Abundance of metabolites in amino sugar and nucleotide sugar metabolism and fructose and mannose metabolism in normal mouse lung tissues (normal) and KL tumor tissues. Individual data points are shown with mean values and SD for six normal and ten KL tissues. **d,** Abundance of metabolites in amino sugar and nucleotide sugar metabolism and fructose and mannose metabolism in normal human lung tissues (normal) and KL tumor tissues. Individual data points are shown with mean values and SD for 18 normal sections and 15 KL tumor sections used in the previous paper^14^. **e,** Enriched KEGG pathways were identified using GSEA against C2 pathways of the MSig database^42^ in the KL mouse tumors compared with matched normal tissues. The top three enriched pathways from the analysis are represented. Their leading-edge genes are shown in Supplementary Fig. 2c and the complete list (FDR q-val <0.05) is available in Supplementary Table 3. Statistical significance was assessed using two-tailed Student’s t-test (c) and (d). **p<0.01; **** p<0.0001. Metabolomics was performed once.

In accordance with results from our previous study on human NSCLC cell lines and tumors^14^, KL co-mutant tumors had hallmarks of altered metabolism of nitrogen-related pathways; in the current analysis of murine tumors, these included pathways involving amino acids, pyrimidines and amino sugars/nucleotide sugars (Fig. 1b and Supplementary Table 2). Direct inspection of individual metabolites related to amino sugars/nucleotide sugar pathways revealed that several were elevated in KL co-mutant tumors (Fig.1c and Supplementary Fig. 1c). Primary human KL co-mutant NSCLCs also accumulated amino sugar metabolites compared to adjacent, non-tumor involved lung tissue (Fig. 1d). Gene sets involved in amino sugar and nucleotide sugar metabolism, fructose and mannose metabolism, and oxidative phosphorylation were elevated in KL tumors from mice compared with normal lungs (Supplementary Fig. 2a, b). These gene sets, along with genes related to O-glycan biosynthesis, associated with KL tumors when compared with tumors containing mutations in Kras but not Lkb1 (Fig. 1e, Supplementary Fig. 2c and Supplementary Tables 3-5). It is worth noting that although many metabolites discriminating between normal lungs and co-mutant tumors were shared between humans and mice, mice unlike humans do not induce expression of *CPS1* mRNA (Supplementary Fig. 3a-c)^14^, indicating incomplete cross-species conservation of metabolic adaptations in tumors with these mutations. Although sphingolipid metabolism ranks as a highly altered pathway by metabolomics (Fig. 1b), none of the sphingolipid-related gene sets were associated with KL status (Supplementary Table 4). Because results from both metabolomics and gene set enrichment analysis (GSEA) indicate the association of the ‘amino sugar and nucleotide sugar pathway’ and ‘fructose and mannose metabolism pathway’ with KL co-mutation status, we focused on how these two pathways impact tumors with KL co-mutation.

### The hexosamine biosynthesis pathway is upregulated in KL co-mutants

Amino sugar/nucleotide sugar metabolism and fructose and mannose metabolism all contribute to the hexosamine biosynthesis pathway (HBP), which uses glycolytic intermediates (fructose-6-phosphate, F6P), uridine, acetyl-CoA and glutamine (Fig. 2a and Supplementary Fig. 4a). The first and rate-limiting step is transfer of the amide nitrogen from glutamine to fructose-6-phosphate by glutamine-fructose-6-phosphate transaminase (GFPT) to generate glucosamine-6-phosphate (GlcN-6-P). GlcN-6-P is acetylated to become N-acetyl glucosamine 6-P (GlcNAc-6-P) and then converted to N-acetyl glucosamine 1-P (GlcNAc-1-P) via a mutase reaction. Finally, it reacts with UTP, forming UDP-N-acetylglucosamine (UDP-GlcNAc), the end product of this pathway^15^.

**Figure 2.**
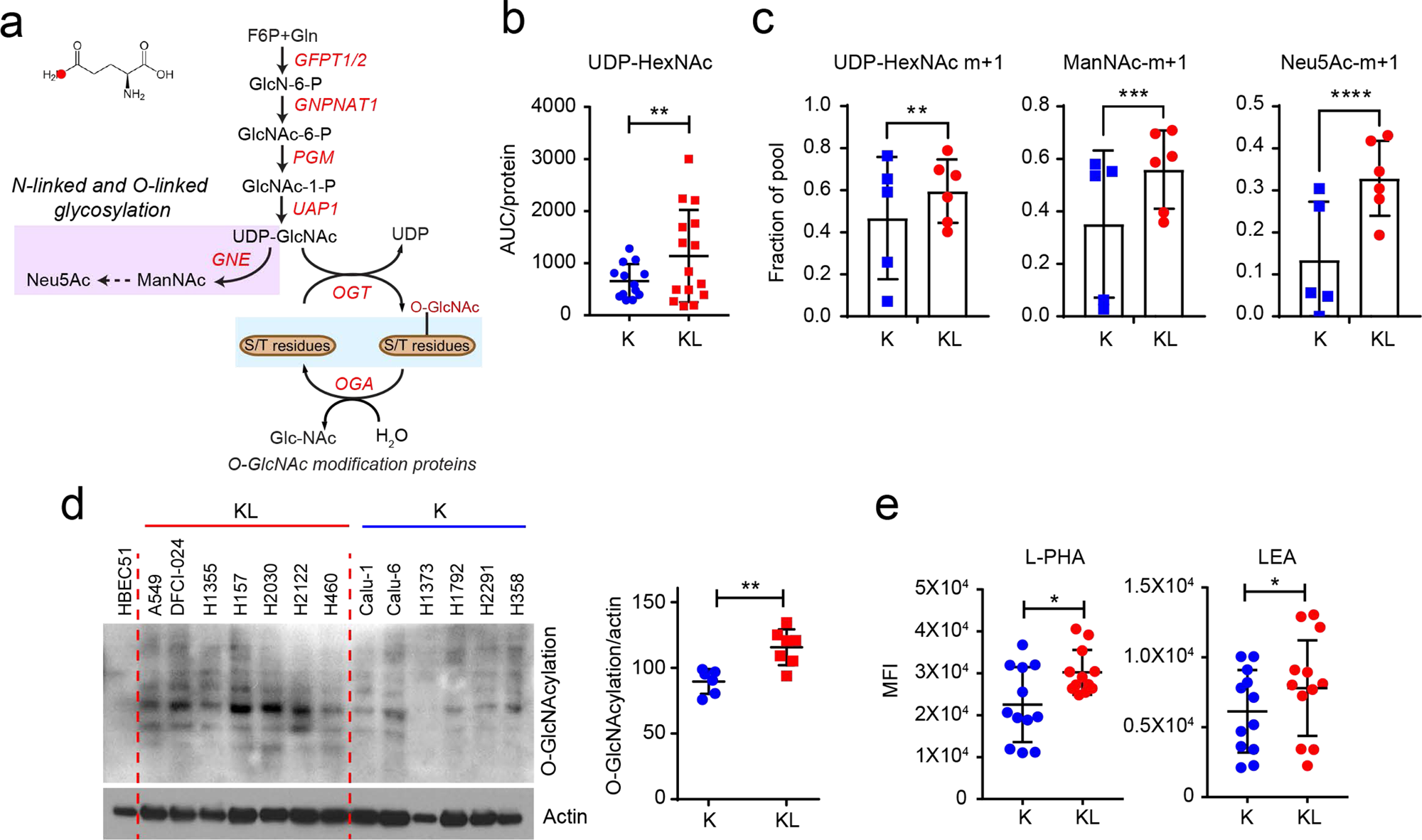
The hexosamine biosynthesis pathway is upregulated in KL cells. **a,** Schematic of the hexosamine biosynthesis pathway (HBP). Metabolites in the glycosylation pathway are in lilac and O-GlcNAcylation is in light blue. Schematic of ^15^N incorporation from [γ-^15^N]glutamine into the HBP intermediates is shown in Supplementary Fig. 4a. **b,** Abundance of UDP-HexNAc in K and KL cell lines. AUC= area under the curve. Individual data points represent replicates from each cell line and are shown with mean values and SD for five K and KL cell lines. Three replicates were analyzed for each cell line, except for H441 and H358, for which two replicates were analyzed. **c,** ^15^N labeling in UDP-HexNAc, ManNAc, and Neu5Ac in K and KL cells cultured with [γ-^15^N]glutamine containing media for 6 hours. Individual data points represent the average value of each cell line (3 replicates/cell line). **d,** *Left*, Global O-GlcNAcylation in K and KL cells. Human bronchial epithelial cell line HBEC51 is used as control. *Right*, Quantification of O-GlcNAcylation normalized by actin levels. **e,** Cell surface L-PHA and LEA lectin binding in K and KL cells was measured by flow cytometry. Statistical significance was assessed using two-tailed Student’s t-test (b), (c), (d), and (e). In **c,** statistical analysis was done with individual replicate of each cell lines (n=3/each cell line). *p<0.05; **p<0.01; ***p < 0.001; ****p<0.0001. In targeted metabolomics, single time point stable isotope labeling for ManNAc and Neu5Ac was performed once, and single time point stable isotope labeling for UDP-HexNAc was performed twice. Western blots were repeated three times or more. FACS analyses were performed twice.

To examine the HBP in human NSCLC cells, we measured UDP-HexNAc as a surrogate of UDP-GlcNAc. UDP-GlcNAc is not separated from UDP-GlaNAc in our LC/MS system, although the UDP-HexNAc peak is almost entirely composed of UDP-GlcNAc^16-18^ in a panel of cell lines with either KRAS (K) or KRAS/LKB1 (KL) mutations. Steady-state levels of UDP-HexNAc were modestly elevated in KL cells (Fig. 2b). The amide nitrogen donated by glutamine at the GFPT step is transmitted to downstream amino sugar metabolites including N-acetylmannosamine (ManNAc) and N-acetylneuraminic acid (Neu5Ac) (Supplementary Fig. 4a). To evaluate HBP flow, we measured transfer of ^15^N from [amide-^15^N]glutamine ([γ-^15^N]glutamine) to UDP-HexNAc. After 6hr of incubation with [γ-^15^N]glutamine, KL cells as a group had uniformly high labeling in HBP intermediates, whereas labeling in K cell lines was heterogenous (Fig. 2c).

UDP-GlcNAc is a substrate for post-translational protein modification by glycosylation and the synthesis of glycolipids, proteoglycans, and glycosylphosphatidylinositol anchors (Fig. 2a and Supplementary Fig. 4a). One of the most common modifications is O-GlcNAcylation of cytoplasmic, nuclear and mitochondrial proteins to regulate signaling, transcription and other functions^19^. In our cell line panel, KL cells had higher levels of global O-GlcNAcylation than K cells (Fig. 2d). UDP-GlcNAc is also required for O-linked and N-linked glycosylation of membrane proteins, especially for both initiation of N-glycosylation in the endoplasmic reticulum (ER) and N-glycan branching and antenna elongation in the Golgi apparatus^20,21^. GlcNAc-branched N-glycans are recognized by the lectin phytohemagglutinin-L (L-PHA)^22^, while poly N-acetyllactosamine (polyLacNAc) extension of N-linked glycan antennae is recognized by the lectin *Lycopersicon esculentum* (LEA). To compare N-linked glycan structures between K and KL cells, we measured surface binding of L-PHA and LEA by FACS analysis. KL cells displayed enhanced binding of both L-PHA and LEA (Fig. 2e), indicating more complex glycan structures on their surface glycoproteins. Taken together, these data suggest that enhanced hexosamine biosynthesis in KL-comutant cancer cells increase both intracellular O-GlcNAcylation and complex N-Glycan structures on cell surface glycoproteins^23^, which may contribute to tumor aggressiveness.

### LKB1 regulates the hexosamine biosynthesis rate

We next examined whether LKB1 loss regulates hexosamine metabolism in KRAS-mutant NSCLC cells. To this end, we engineered two KL cell lines (H460 and H2122) to express wild-type LKB1 (Fig. 3a). Expression of LKB1 reduced the transfer of ^15^N from [γ-^15^N]glutamine into HBP metabolites including GlcNAc-6-P, UDP-HexNAc, and ManNAc (Fig. 3b). Time course labeling with [γ-^15^N]glutamine also demonstrated delayed hexosamine biosynthesis (Fig. 3c). Reduced hexosamine biosynthesis in LKB1-expressing cells led to decreased protein O-GlcNAcylation and decreased complex N-glycan structures on cell surface glycoproteins (Figs. 3d-f and Supplementary Fig. 4b). Collectively, these data demonstrate that LKB1 suppresses protein glycosylation through mechanisms involving reduced HBP synthesis.

**Figure 3.**
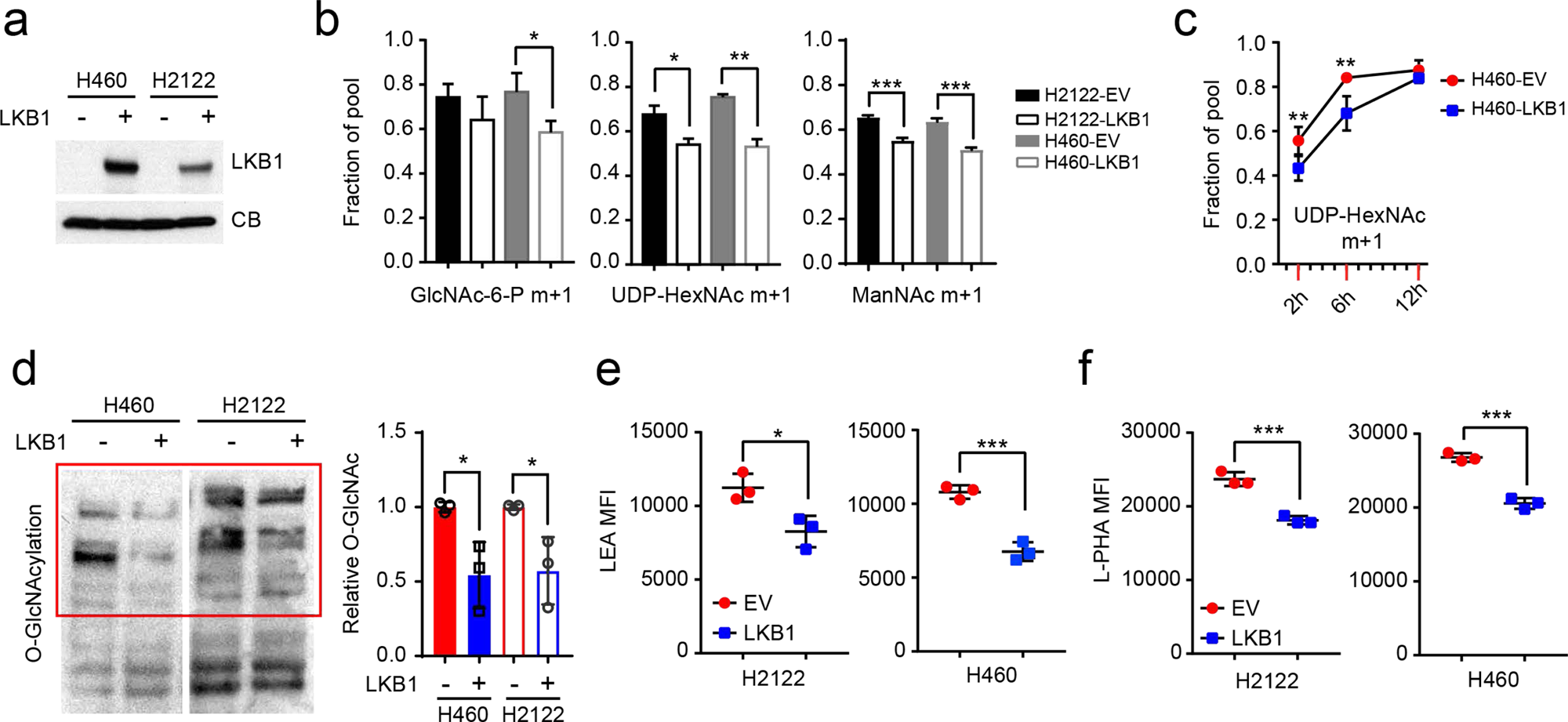
LKB1 regulates the hexosamine biosynthesis pathway. **a,** LKB1 re-expression in H460 and H2122, two KL cells. Cyclophilin B (CB) is used as loading control. **b,** ^15^N labeling in GlcNAc-6-P, UDP-HexNAc and ManNAc in empty vector (EV) control and wildtype LKB1 (LKB1) expressing H460 and H2122 cells cultured with [γ-^15^N]glutamine. **c,** Time-course of ^15^N labeling in UDP-HexNAc in EV and LKB1 expressing H460 cells cultured with [γ-^15^N]glutamine. **d,** *Left*, Global O-GlcNAcylation in EV and LKB1 expressing H460 and H2122 cells. *Right*, Relative intensity of O-GlcNAcylation (in red bracket) to a loading control (cyclophilin B, CB). **e,** Cell surface LEA lectin binding in EV and LKB1-expressing H460 and H2122 cells was measured by flow cytometry. **f,** Cell surface L-PHA lectin binding in EV and LKB1 expressing H460 and H2122 cells was measured by flow cytometry. Data in **b** and **c** are the average and SD of three independent cultures. In **b, e, f,** statistical significance was assessed using two-tailed Student’s t-test. In **c,** to calculate significance on repeated measurements over time, two-way ANOVA was used. Stable isotope experiments and O-GlcNAcylation western blotting were repeated twice. All other experiments were repeated three times or more. *p<0.05;**p<0.01; ***p<0.005.

### KL cells are selectively vulnerable to GFPT inhibition

To test whether enhanced hexosamine biosynthesis imposes metabolic liabilities, several K and KL cell lines were treated with inhibitors of hexosamine biosynthesis and N-linked glycosylation, including azaserine (GFPT inhibitor), OSMI-1 (O-GlcNAc transferase (OGT) inhibitor) and siRNA against OGT (Fig.4a). Inhibiting either GFPT or OGT more potently suppressed growth of KL cells (Fig. 4b-d and Supplementary Fig. 5a-f) although efficiency of OGT silencing or of OSMI-1 treatment was comparable between K and KL cells (Supplementary Fig. 5c,d and f), indicating that KL cells are more dependent on the HBP for survival. Azaserine treatment also markedly enhanced cell death, but only in KL cells (Supplementary Fig. 6a-c). Of note, in all four KL cell lines tested, azaserine’s toxicity at 0.5µM (the average IC50 among KL cells) was at least partially reversed by supplementing with N-acetyl glucosamine (GlcNAc), a metabolite that enters the HBP at GlcNAc-6-P after phosphorylation by N-acetylglucosamine kinase (NAGK) (Fig. 4a,e). This finding indicates that in these cells, azaserine’s mechanism of action involves HBP suppression. This is important because azaserine inhibits a number of amidotransferases and other enzymes in addition to GFPT. Collectively, these data show that KL cells require GFPT and the HBP for survival, and do so to a greater degree than K cells.

**Figure 4.**
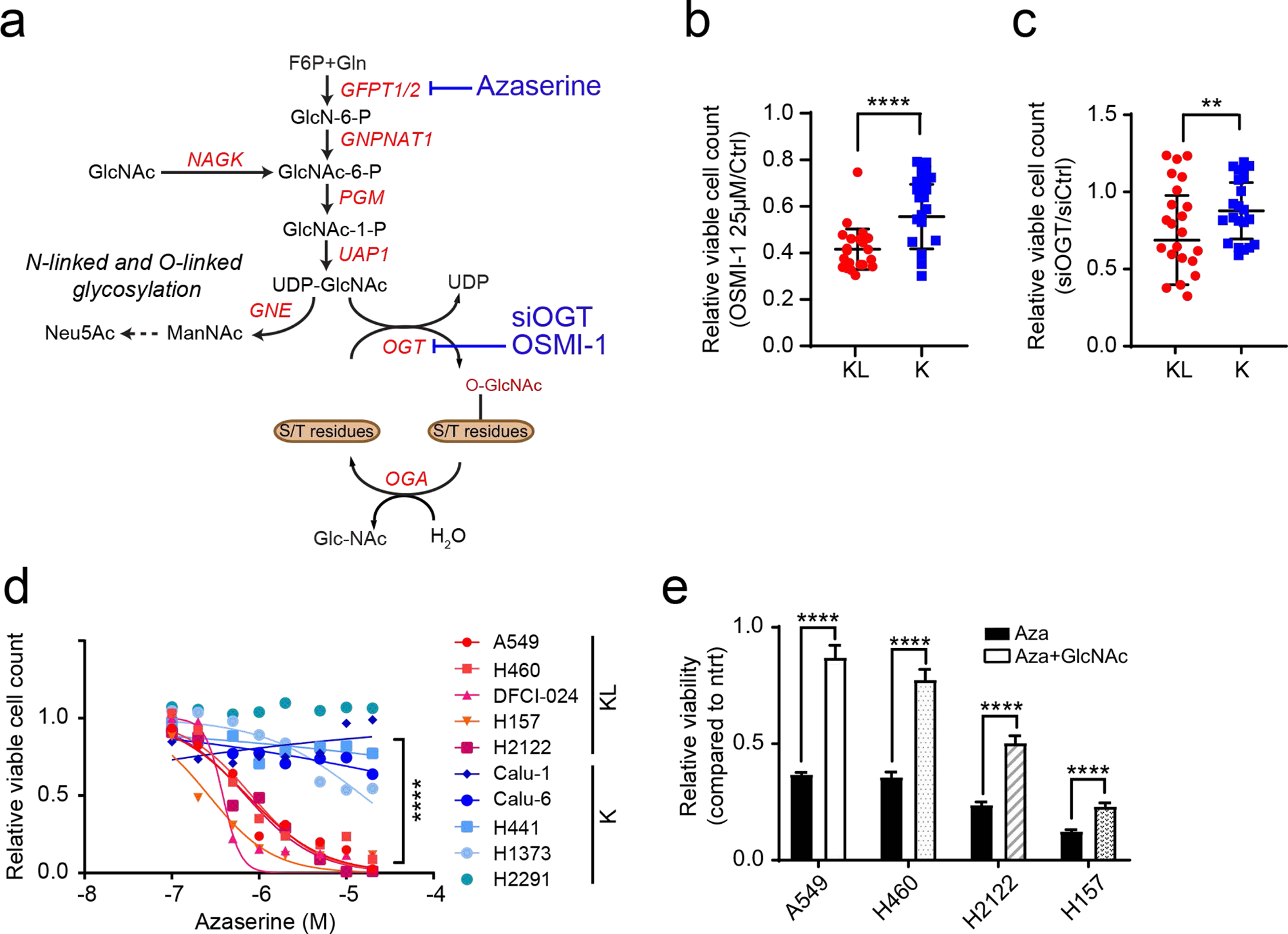
KL cells are dependent on the rate limiting step of the hexosamine biosynthesis pathway. **a,** Schematic of the hexosamine biosynthesis pathway. Enzyme/pathway targeted by each inhibitor and siRNA is shown. **b,** Relative viability of a panel of K and KL lines following a 72hr exposure to OSMI-1 (25µM). **c,** Relative viability of K and KL lines to OGT silencing for 96hr. **d,** Dose-response curves for K and KL lines following 72hr exposure to azaserine. **e,** Effect of GlcNAc (40mM) on azaserine (0.5µM)-treated KL cells (four cell lines, n=6/line). Data are average and SD of three or more independent cultures. Statistical significance was assessed using unpaired t-test with Welch’s correction (**b, c**), Mann-Whitney U test (**d**), two-tailed Student’s t-tests (**e**); **p<0.01; **\*\***\*\*p<0.0001. Cell viability assay of OSMI-1 and dose-response assay were performed three times or more. OGT silencing assay and viability assay with azaserine treatment in the presence or absence of GlcNAc were repeated twice.

### KL cells and tumors require GFPT2

Two GFPT isoforms, GFPT1 and 2, are encoded by separate genes with different tissue distributions^24^, and either or both of these isoforms could account for azaserine toxicity in KL cells. To identify the target of azaserine in NSCLC cells, we transfected K and KL cells with siRNAs targeting either GFPT1 or GFPT2. Only H2122 was affected by GFPT1 silencing (Fig. 5a and Supplementary Fig. 7a). On the other hand, silencing GFPT2 reduced viability in all KL cell lines tested, but no K cells (Fig.5b and Supplementary Fig. 7b). Supplementing GlcNAc completely reversed the effects of GFPT2 silencing in H460 KL cells (Fig. 5c). High levels of *GFPT2* mRNA, but not *GFPT1* mRNA, correlate with poor prognosis in NSCLC cohorts (Supplementary Fig. 7c,d). GFPT2 protein levels tend to be higher in KL cells while GFPT1 levels are invariable between K and KL cells (Supplementary Fig.7e,f). H2122, the only cell line to respond to GFPT1 silencing, had the lowest expression of GFPT2 protein. We generated two KL cell lines (H460 and H157) with doxycycline (Dox)-inducible expression of a CRISPR-Cas9 KO system where Cas9 is induced by Dox treatment^25^ and GFPT2 becomes depleted (Supplementary Fig. 7h). While Dox-mediated GFPT2 depletion barely affected growth in a monolayer culture, it suppressed anchorage-independent cell growth and invasion capacity (Fig. 5d,e and Supplementary Fig. 7g). GFPT2 KO clones generated by conventional, constitutive CRISPR-mediated genome editing^26^ decreased both proliferation and colony formation (Fig. 5f,g and Supplementary Fig. 7i,j). Supplementing GlcNAc normalized colony formation and O-GlcNAcylation in GFPT2 KO cells (Supplementary Fig. 7k,l), as expected if the GFPT2 CRISPR effects are through HBP suppression. Consistent with this, GFPT2 deletion reduces intracellular levels of HBP intermediates and N-glycan branching of glycoproteins on the cell surface (Fig. 5h,i and Supplementary Fig. 7m,n). KL cells reconstituted with LKB1 lost sensitivity to GFPT2 silencing, indicating that dependence on GFPT2 in KL cells is mediated by LKB1 (Fig. 5j).

**Figure 5.**
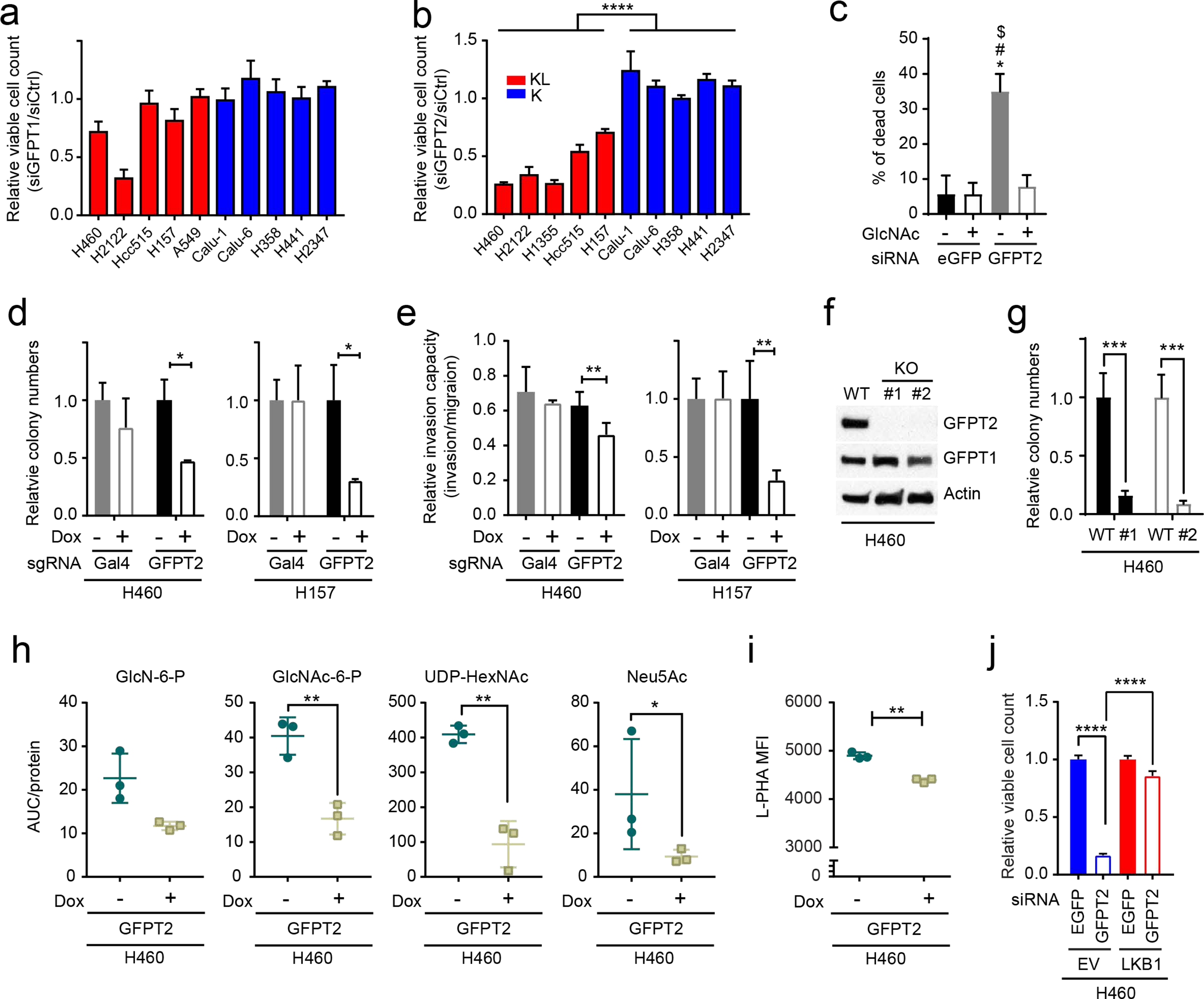
KL cells require GFPT2. **a and b,** Sensitivity to GFPT1 (**a**) and GFPT2 (**b**) silencing in K and KL cells (n = 6). **c,** Rescue effect of GlcNAc supplementation on GFPT2 silencing-induced cell death (n=6). **d,** Effects of a Dox-induced GFPT2 sgRNA (GFPT2) on anchorage independent growth of H460 and H157 (n = 3 for H157, n=4 for H460). Gal4 is a Dox-inducible control sgRNA. **e,** Effects of Dox-induced GFPT2 KO on invasion capacity of H460 and H157 cells (n=4 for H460-sgGal4-/+ Dox, n=8 for H460-sgGFPT2-/+Dox, n=12 for H157 GFPT2-Dox and H157-Gal4-Dox, n=8 for H157 GFPT2+Dox, n=7 for H157-Gal4+Dox). **f,** Abundance of GFPT1 and 2 in parental and GFPT2 KO H460 cells. Actin was used as a loading control. **g,** Effects of GFPT2 KO (two clones) on anchorage independent growth of H460. **h,** Abundance of hexosamine metabolites in Dox-inducible GFPT2 KO H460 cells (n=3). sgGal4 is used as a Dox-inducible sgRNA control. **i,** Effect of Dox-inducible GFPT2 KO H460 cells on cell surface L-PHA lectin binding. **j,** Effect of LKB1 on GFPT2 silencing-induced loss of viability (n=5). In **b, d, e, g, h, and i,** statistical significance was assessed using two-tailed Student’s t-test. In **c** and **j,** statistical significance was assessed using one-way ANOVA followed by Tukey’s post hoc test was used. In **c,** *, p<0.05 comparing to sieGFP without GlcNAc; #, p<0.05 comparing to sieGFP with GlcNAc treatment; $ p<0.05 comparing to siGFPT2 with GlcNAc treatment. *p<0.05; **p<0.01; ****p<0.0001. Western blot was repeated three times and all other experiments were performed twice.

Next, to examine the effect of GFPT inhibition on tumor growth and in vivo HBP suppression, nude mice were subcutaneously injected with two KL (A549, H460) and two K cell lines (Calu-1, Calu-6) and treated with azaserine when palpable tumors were present. Compared with vehicle control, azaserine attenuated growth of both KL tumor lines, but neither K tumor line (Fig. 6a and Supplementary Fig. 8a,b). We then examined turnover of HBP metabolites by infusing mice bearing subcutaneous A549 xenografts with [γ-^15^N]glutamine after two weeks of vehicle control or azaserine treatment. Tumors from the azaserine-treated mice displayed lower incorporation of ^15^N into GlcNAc-6-P and UDP-HexNAc (m+1 labeling) compared with those from control mice (Fig. 6b). Consistent with the in vivo infusions, targeted metabolomics of two KL tumors revealed depletion of several HBP intermediates in azaserine-treated tumors (Fig. 6c).

**Figure 6.**
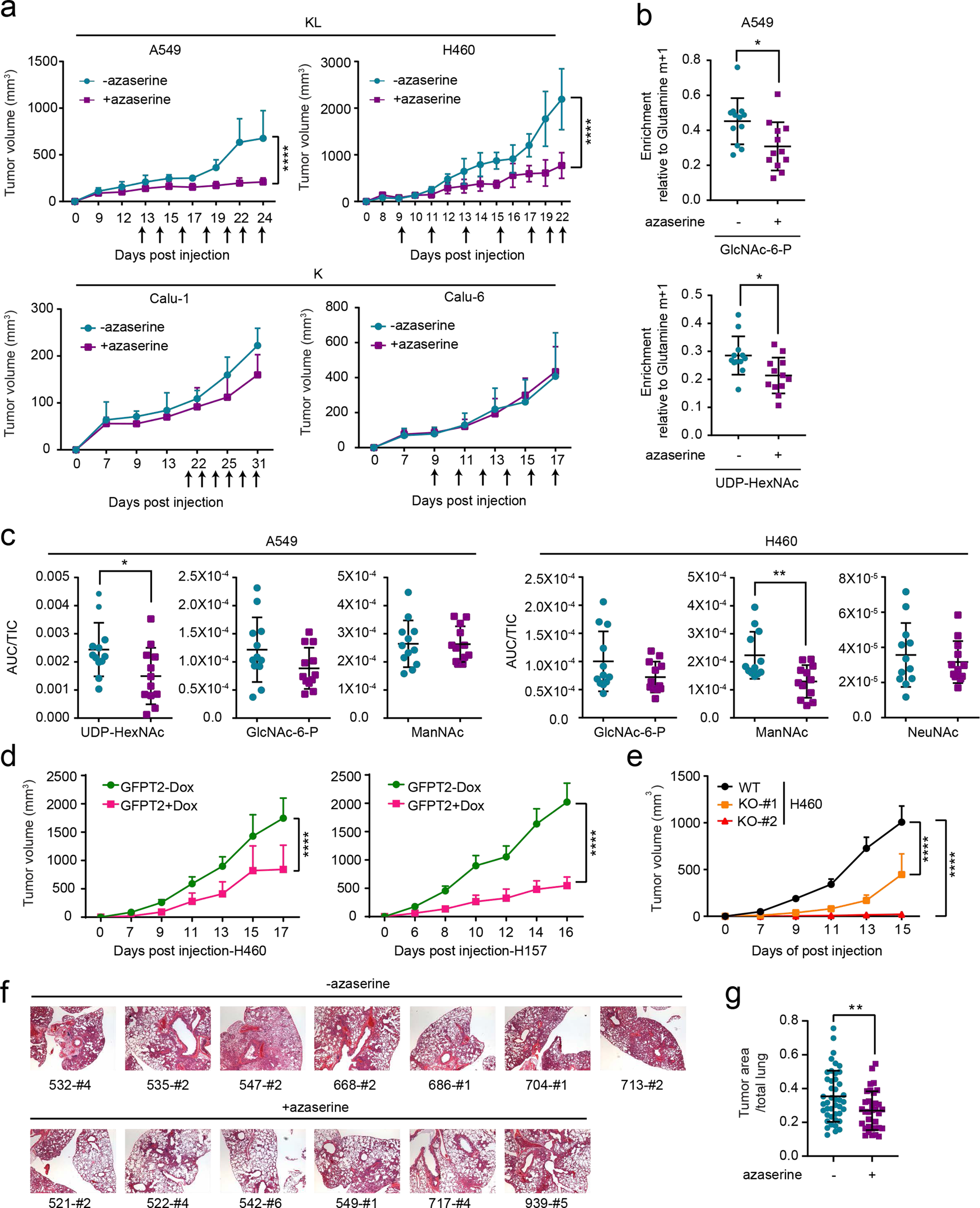
GFPT2 suppression inhibits KL tumor growth. **a,** *Top*, growth of A549 (left) and H460 (right) xenografts in presence and absence of azaserine (2.5mg/kg, qod for 6-7 times total, arrows indicate when azaserine was injected). *Bottom*, growth of Calu-1 (left) and Calu-6 (right) xenograft in presence and absence of azaserine. Mean tumor volume and SD are shown for each group (n=4 for A549, n=4 for H460, n=5 for both Calu-1 and Calu-6. Combined results from two independent H460 xenograft experiments (total n=8) are shown here). **b,** ^15^N labeling in GlcNAc-6-P (top) and UDP-HexNAc (bottom) in mice bearing A549 treated (purple) or non-treated (turquoise) with azaserine. **c,** Abundance of hexosamine metabolites in A549 (first three panels) and H460 (last three panels) xenografts in Fig. 6a. AUC/TIC=Area under the curve/total ion count. Individual data points are shown with mean values and SD for 12 sections (three fragments per tumor). **d,** Growth of Dox-inducible GFPT2 KO H460 (left) and H157 (right) xenografts in presence and absence of Dox. Mean tumor volume and SEM are shown for each group (n=5 per group). **e,** Growth of H460 WT or GFPT2 KO (two different clones) xenografts. Mean tumor volume and SEM are shown for each group (n=5 per group). **f,** Representative images of non-treated KL mice (upper panel, total 7 mice) and azaserine treated mice (lower panel, total 6 mice). Numbering is [mouse identification]-[image #]. **g,** Tumor area from Fig. 6f was quantified with imageJ and % of tumor burden out of total lung was analyzed. 39 images from non-treated mice and 34 images from azaserine-treated mice were used for quantification. Significance was assessed using two-tailed Student’s t-tests (**b, c, g**); two-way ANOVA with Sidak’s multiple comparisons test (**a, d**); two-way ANOVA with Tukey’s multiple comparisons (**e**). *p<0.05; **p<0.01; ****p<0.0001. Tumor growth studies in **e** and **f,** in vivo infusion in **b,** and targeted metabolomics in **c** were performed once. Tumor growth study in **a** and **d** were repeated twice.

We also assessed the effect of conditional GFPT2 CRISPR on tumor growth. Nude mice were injected subcutaneously with two KL cell lines (H460, H157) and one K cell line (Calu-1) expressing sgGFPT2 and treated with or without Dox. Dox induction of Cas9-mediated GFPT2 KO reduced tumor growth in H460 and H157, but not Calu-1 (Fig. 6d and Supplementary Fig. 8c-e). Tumor growth suppression was also observed after constitutive CRISPR-mediated GFPT2 loss in H460 cells (Fig. 6e and Supplementary Fig. 8f).

Finally, to assess the efficacy of HBP suppression in genetically engineered mouse models with autochthonous NSCLC and an intact immune system, we used mice with a genotype of Kras^G12D/wt^/Lkb1^LoxP/LoxP1^. At 4-5 weeks after inhalation of an Ad5-CMV-Cre, when tumor burden reaches ∼5% of the total pulmonary volume^27,28^, the mice were randomized to receive vehicle or azaserine (2.5mg/kg every other day, 6-7 times total). Mice were euthanized after the final treatment and lung tumor tissues were analyzed histologically. Azaserine monotherapy reduced tumor burden relative to total lung volume (Fig. 6f,g).

## Discussion

The simultaneous incidence of oncogenic KRAS and loss of LKB1 has been shown to specify aggressive oncological behavior both in mouse models of lung cancer and human patients. In this study, we provide evidence that KL tumor aggressiveness is associated with dependence on the HBP through GFPT2 (Supplementary Fig. 9). LKB1 loss in the context of oncogenic KRAS plays a prominent role in the flux of glutamine into the HBP through GFPT2, the rate-limiting step of the HBP. GFPT also feeds the glutamate pool, which may complement glutaminase’s contribution to anaplerosis. In pancreatic ductal adenocarcinoma (PDAC), oncogenic KRAS mutation stimulates glucose-dependent biosynthetic pathways including the HBP^6^. Ras oncogenes have been shown to induce N-glycosylation and oncogenic KRAS is associated with complex N-glycan structures in colorectal cancer cells^29,30^. In line with these previous findings, our GSEA of metabolic gene transcript profiles from normal lung tissues versus KRAS mutant lung tumors also returned “fructose and mannose metabolism,” a gene set related to the HBP, as highly associated with KRAS mutation status. When we compared KL co-mutant lung tumors with KRAS tumors, however, the association of genes in HBP-related pathways was even stronger in the co-mutants. This is consistent with the elevated rate of ^15^N entry into HBP metabolites in KL co-mutant cells, suppression of this activity by LKB1, and increased sensitivity to GFPT inhibition in commutant cells and tumors. Thus, our data collectively suggest that, at least in NSCLC, LKB1 loss in the context of oncogenic KRAS alters metabolic preferences, rendering cancer cells dependent on the HBP and subsequent glycosylation pathways.

Another difference between PDAC and NSCLC is related to isoform specificity of HBP dependence. GFPT2 and GFPT1 both catalyze the entry step into the HBP. In PDAC, GFPT1 is required for glycosylation and KRAS regulates GFPT1 expression^31^. In our study, however, GFPT1 silencing barely impacted NSCLC cell survival regardless of their mutation status, and expression of *GFPT2* but not *GFPT1* correlated with poor clinical outcomes in LUAD. The reason for this isoform specificity is unknown, but may involve its association with the epithelial-mesenchymal transition (EMT). EMT transcription factors *SNAI1* and *TWIST1* are highly correlated with *GFPT2*, but not *GFPT1*, in clinical lung adenocarcinoma samples^32^. Upregulation of *GFPT2* expression was observed in transcriptomic signatures of both mesenchymal lung tumors in mice and the mesenchymal, or claudin-low, subtype of triple-negative breast cancer^13,32^. Given that KL co-mutants emulate claudin-low breast cancer, which are aggressively metastatic tumors enriched with self-renewing tumor-initiating cells^33^, and that *GFPT2*, not *GFPT1*, is one of the 10 upregulated, claudin-low signature genes found in KL NSCLC^13^, our current findings imply that GFPT2, not GFPT1, is associated with tumor aggressiveness in lung cancer.

We demonstrated that LKB1 loss is required for GFPT2 addiction in KRAS-mutant NSCLC. To fully understand the mechanism by which LKB1 inhibits GFPT2, further work will be required. AMPK is a major metabolic effector and it has been reported to inhibit GFPT1 enzyme activity by phosphorylation^34^. This raises the possibility that AMPK may also be involved in GFPT2 regulation. It will be worthwhile to investigate how GFPT2 activity is perturbed by LKB1, either transcriptionally, translationally, or post-translationally, and which downstream effectors are required to regulate GFPT2.

There are no clinically-approved drugs to inhibit GFPT2 in a well-tolerated manner. Although we found that azaserine selectively reduces viability of KL co-mutant cells, this compound also inhibits purine biosynthesis and other pathways^35^. Prolonged treatment of azaserine (> 3weeks) induces tumor growth in rats^36^. UDP-GlcNAc, a major end product of the HBP, is required for both glycosylation of membrane proteins and O-GlcNAcylation of intracellular proteins. Thus, inhibiting these glyco-functionalization pathways may mimic azaserine treatment. A recent study reported that FR054, a specific inhibitor of the HBP enzyme N-acetylglucosamine-phosphate mutase (PGM3), has an anti-breast cancer effect both in vitro and in vivo and is relatively well tolerated^37^. PGM3 catalyzes the conversion of GlcNAc-6-P into GlcNAc-1-P. Its modulation permits control of both O-GlcNAcylation and glycosylation, and PGM3 inhibition reduces HBP flux and levels of complex N-glycans and O-GlcNAcylation. FR054 treatment may phenocopy GFPT2 suppression and have therapeutic potential for KL co-mutant tumors. It will also be interesting to determine whether HBP inhibition synergizes with other metabolic or cytotoxic therapies in NSCLC.

We previously reported that the urea cycle enzyme carbamoyl phosphate synthetase 1 (CPS1) provides an alternative pool of carbamoyl phosphate to maintain pyrimidine availability and supports survival of KL NSCLC. Our current study demonstrates the significance of GFPT2 in the HBP, another nitrogen-related metabolic pathway, in this aggressive subtype of lung cancer. O-GlcNAcylation, functional consequence of the HBP, can stabilize proteins involved in tumorigenic processes such as c-Myc and β-catenin^38,39^. Glycan modification, another arm of the HBP-mediated protein modification, modulates receptor tyrosine kinases, proteoglycans, cadherins and integrins, which contribute to malignant cells’ ability to invade, migrate and metastasize^40,41^. Upregulation of the HBP in KL co-mutants, therefore, may not only be required for primary cancer growth but also for tumor aggressiveness. The fact that cells with mutation of either KRAS or LKB1 do not require GFPT2 suggests that the metabolic effects mediated by KL co-mutations are essential for GFPT2 addiction. One possible explanation would be that enhanced glucose and glutamine uptake by oncogenic KRAS might provide ample supply for the HBP while LKB1 loss enhances the HBP through GFPT2 (Supplementary Fig. 9). Collectively, our findings suggest the use of GFPT2 or related pathway components as therapeutic targets in KL mutant lung adenocarcinomas, providing a new mechanism of oncogene addiction.

## Acknowledgment

R.J.D. is supported by the Howard Hughes Medical Institute and by grants from the National Cancer Institute (NCI) (R35CA22044901) and the Cancer Prevention and Research Institute of TEXAS (CPRIT) (RP160089). J.K. (UIC) is supported by the NCI (1K22CA226676-01A1), American Lung Association (LCD-614827) and the V Foundation (V2019-022). J.D.M. is supported by the NCI (SPORE P50CA070907) and CPRIT (RP160652). J.K. (UTSW) is supported by NCI (1R01CA196851, SPORE P50CA70907) and the American Cancer Society (RSG-16-090-01-TBG). S.K. was supported by the National Heart, Lung, and Blood Institute (5T32HL098040).

R.J.D. is an advisor for Agios Pharmaceuticals. J.D.M. receives cell line licensing royalties from the NIH and UTSW.

## Supplementary Information

### Legends to Supplementary Data Figures

**Supplementary Data Fig. 1.**
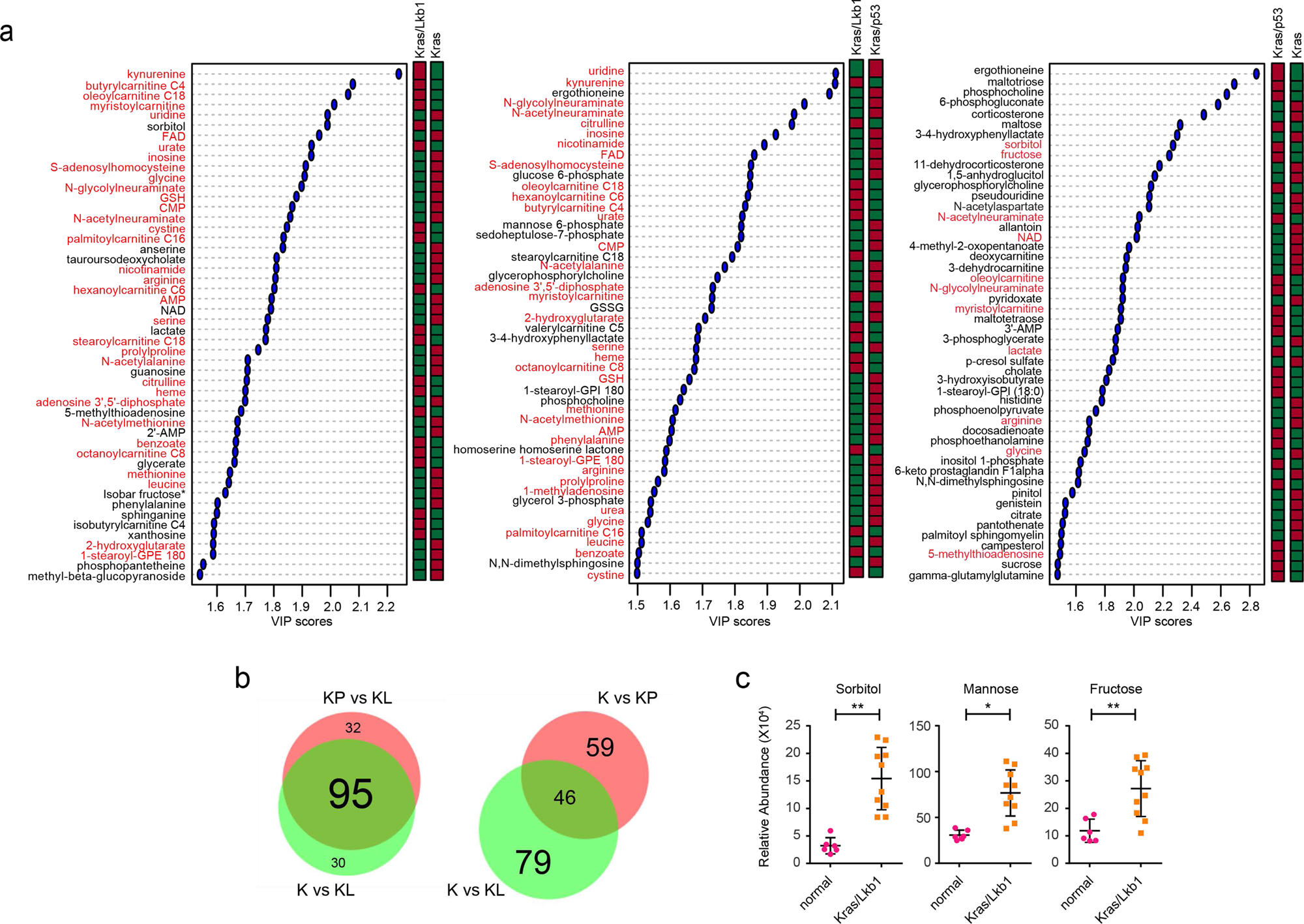
Lkb1 mutation significantly alters metabolism of Kras mutant mouse lung tumor while p53 mutation does not. **a,** Key metabolites differentiating between Kras (K) and Kras/Lkb1 (KL) tumors (left), Kras/Lkb1 (KL) and Kras/p53 (KP) tumors (middle) and Kras (K) and Kras/p53 (KP) tumors (right) (VIP>∼ 1.5). Metabolites shown in all three groups are in red. Relative metabolite abundance is indicated in the bar, with red representing metabolite accumulation. **b,** Venn diagrams of Variable Importance in the Projection metabolites (VIP>1.0) between [K and KL tumor tissues] and [KP and KL tumor tissues] (left) and between [K and KP tumor tissues] and [K and KL tumor tissues]. **c,** Abundance of fructose and mannose pathway intermediates from the metabolomics in Supplementary Data Fig. 1a. Individual data points are shown with mean values and SD for six (normal) or ten (KL) mouse tumors. Statistical significance was assessed using Wilcoxon signed rank test. *p<0.05; **p<0.01.

**Supplementary Data Fig. 2.**
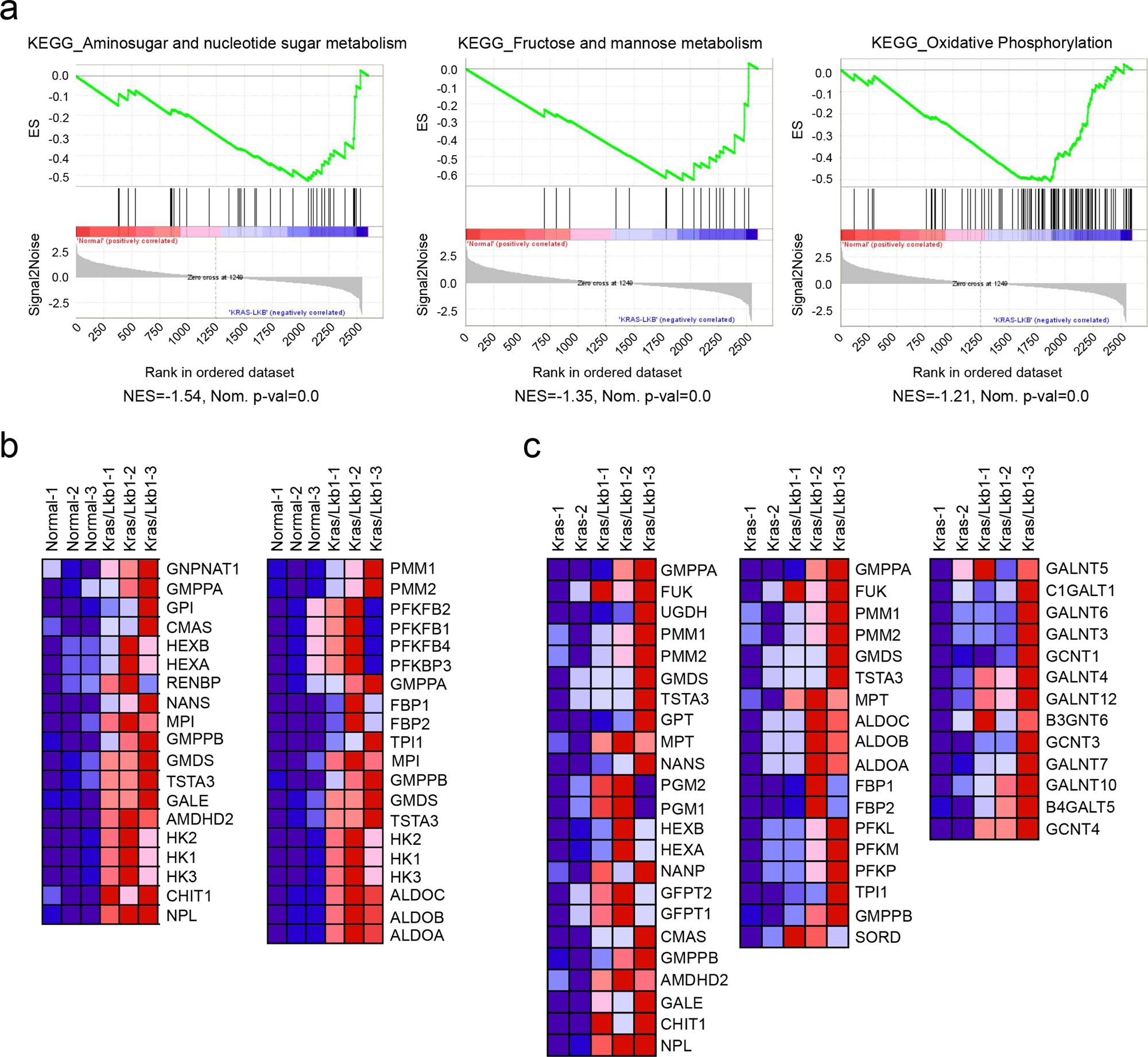
Hexosamine-related metabolic pathways are associated with KL mouse tumors. **a,** GSEA returned ‘‘Amino sugar and nucleotide metabolism”, “Fructose and mannose metabolism”, and “Oxidative Phosphorylation” as the top ranked metabolic gene ontology term from KEGG database. Enrichment statistics include nominal p value and nominal enrichment score (NES). **b,** The leading-edge genes of “Aminosugar and nucleotide sugar metabolism” (left) and “Fructose and mannose metabolism” (right) are in Supplementary Data Fig 2a. The leading-edge genes of “Oxidative Phosphorylation” (not displayed here) are in Supplementary information table 3. **c,** The leading-edge genes of “Aminosugar and nucleotide sugar metabolism”, “Fructose and mannose metabolism” and “O-Glycan biosynthesis” in Fig. 1e.

**Supplementary Data Fig. 3.**
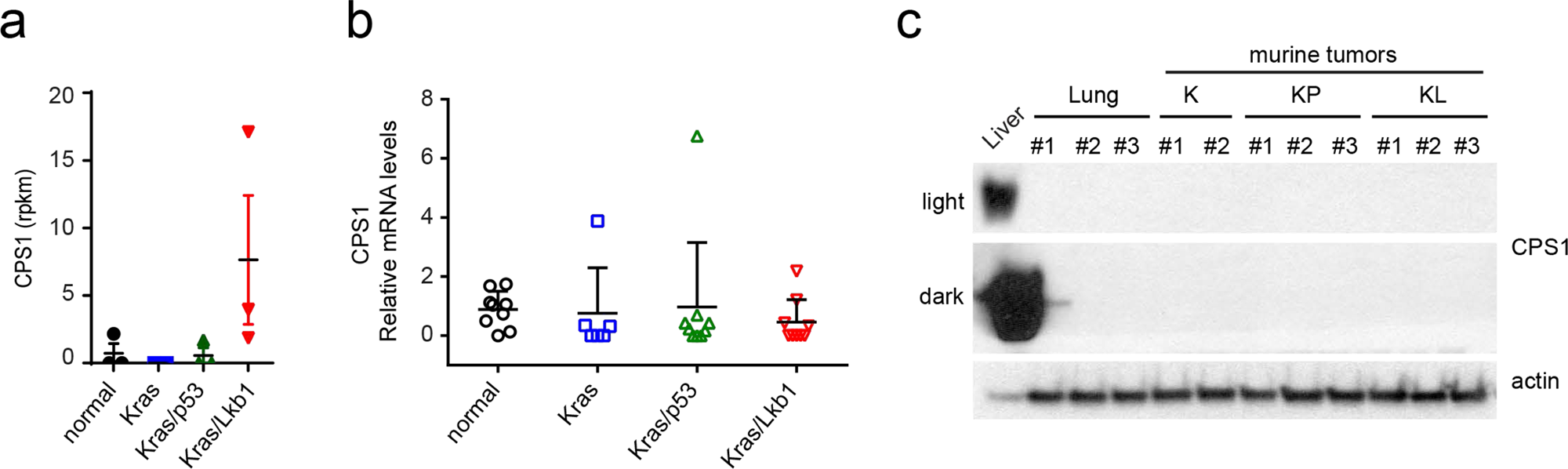
Mouse KL tumors do not display CPS1 induction. **a,** RNAseq result for CPS1 in mouse lung and mouse tumors of the indicated genotypes. Data are the average and SD of tissues from three independent mice except Kras mice (n=2). **b,** CPS1 expression in the tissues used in Supplementary Data Fig. 2a. q-rt-PCR Data are the average and SD of three fragments from three independent mice except Kras mice (three fragments from two mice). **c,** CPS protein expression in the tissues used in Supplementary Fig. 2b. Statistical significance was assessed using one-way ANOVA followed by Tukey’s post hoc test was used and there was no statistical significance among the groups.

**Supplementary Data Fig. 4.**
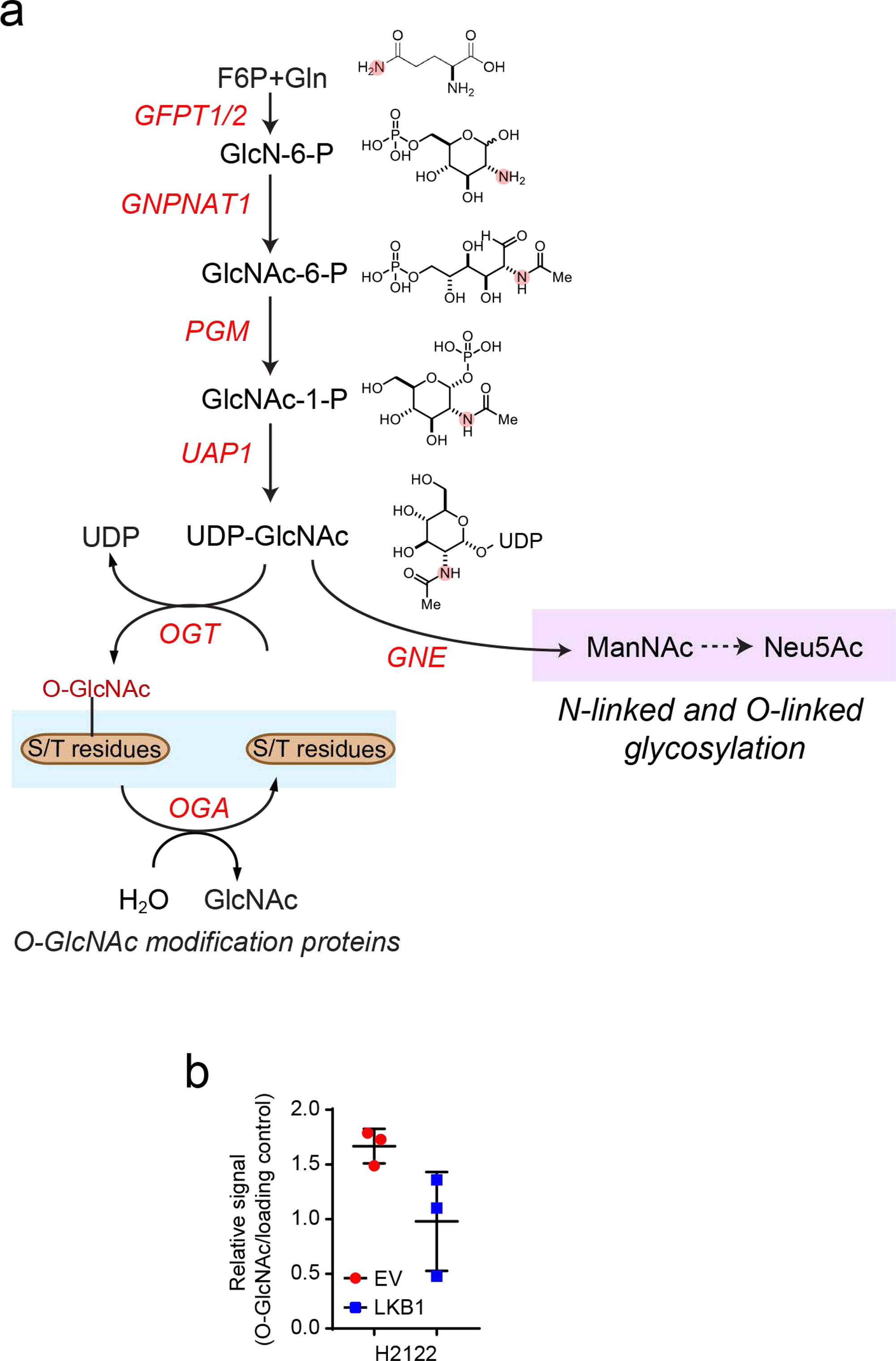
Schematic of the hexosamine biosynthesis pathway and ^15^N incorporation from [γ-^15^N]glutamine (labeled in pink) into the HBP intermediates is shown. **b,** Relative intensity of O-GlcNAcylation to loading control in H2122 cells; O-GlcNAcylation signal was derived from the bracketed region of the blot in Fig. 3d.

**Supplementary Data Fig. 5.**
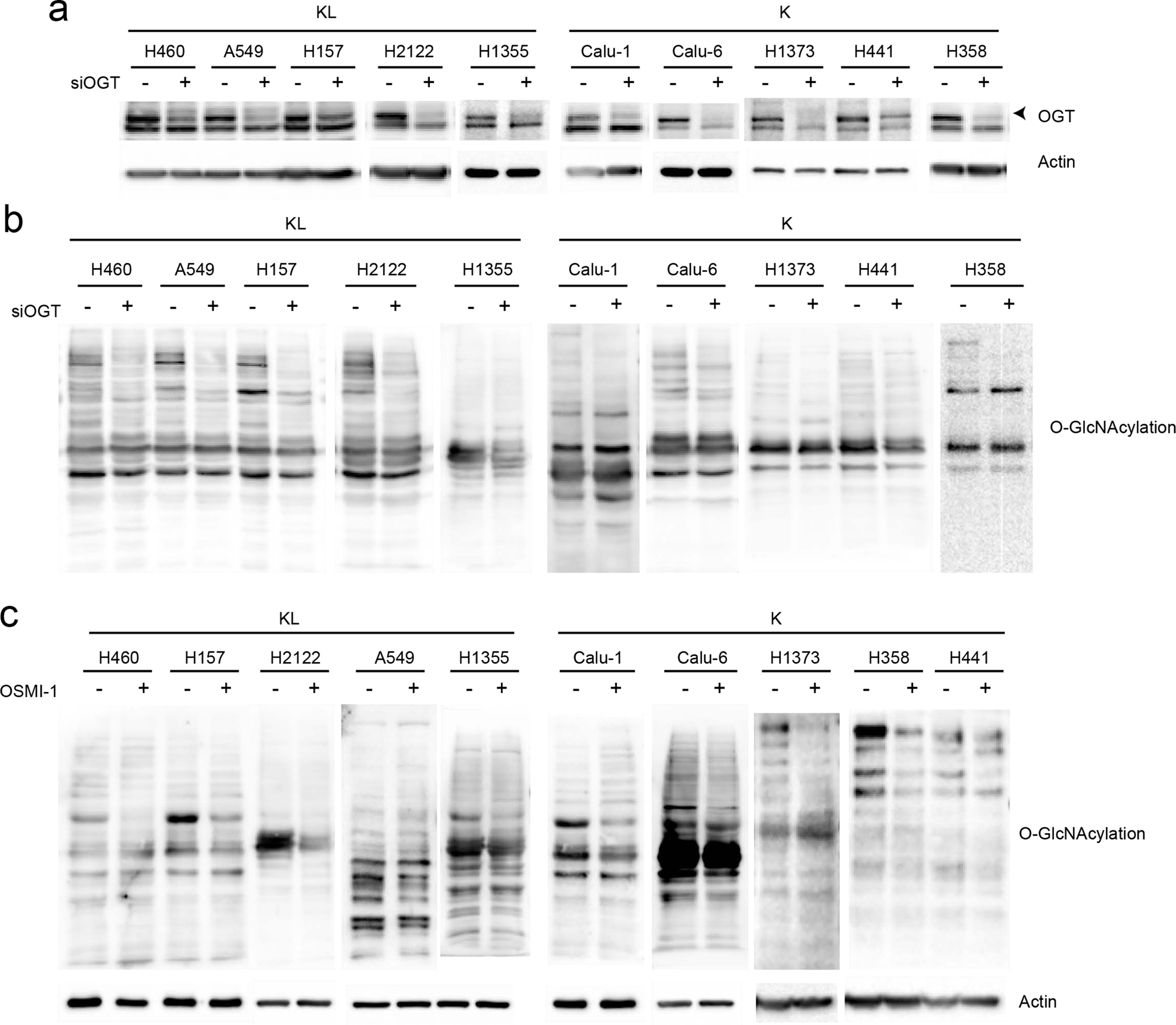
KL cells are sensitive to inhibition of the HBP. **a,** Abundance of OGT protein in K and KL cell lines transfected with a control siRNA or siRNA directed against OGT. Actin is used as a loading control. **b,** Effect of OGT silencing on global O-GlcNAcylation of K and KL cells. **c,** Effect of OSMI-1 treatment on global O-GlcNAcylation of K and KL cells. Actin is used as a loading control. All Western blots are repeated three times or more.

**Supplementary Data Fig. 6.**
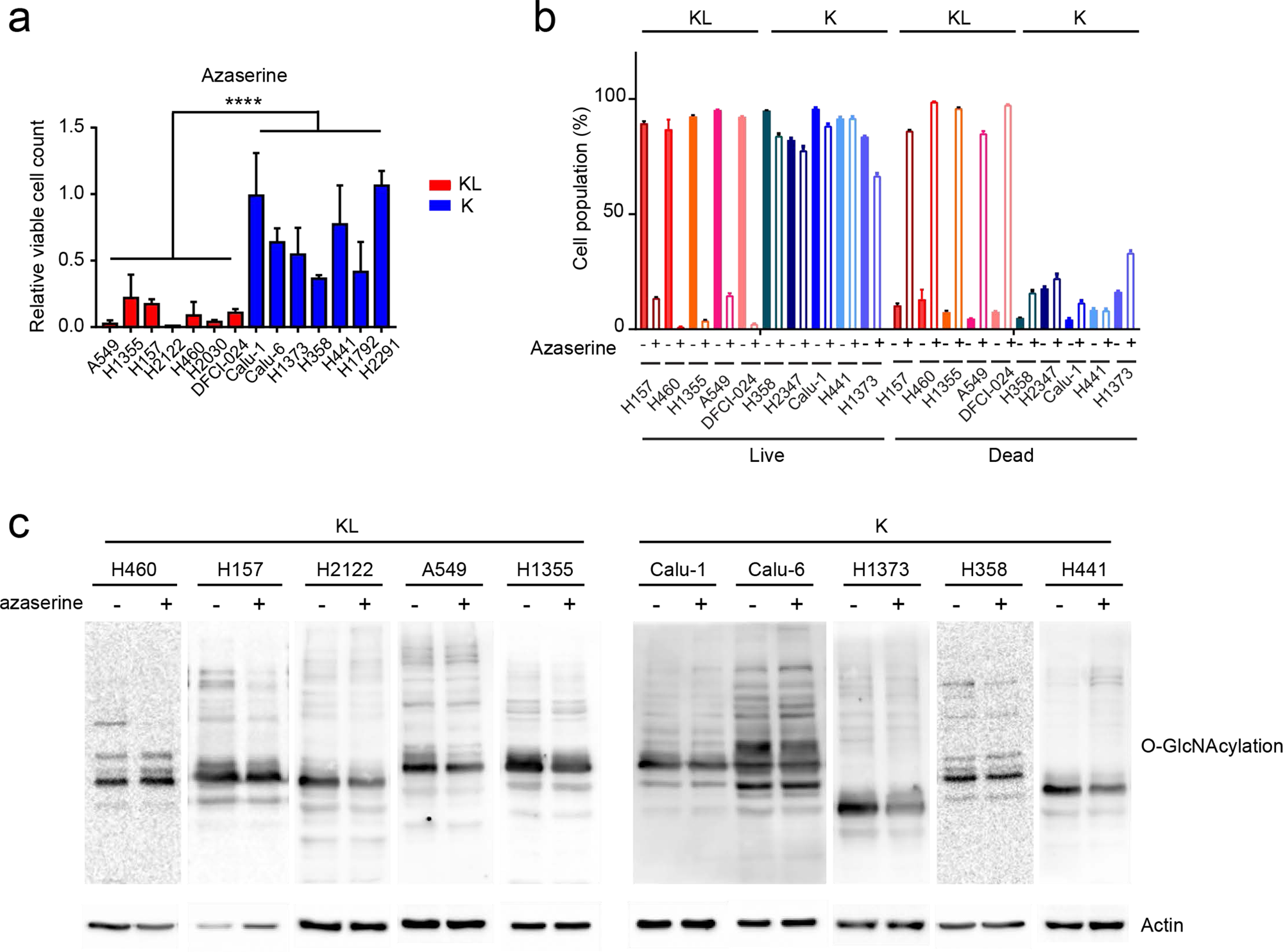
KL cells are sensitive to inhibition of GFPT activity. **a,** Effect of azaserine treatment (1µM) on K and KL cells’ viability (14 cell lines, n=6). Data are the average and SD of six independent cultures. **b,** Effect of azaserine treatment on cell death in K and KL cells (n=3). **c,** Effect of azaserine treatment on global O-GlcNAcylation of K and KL cells. Actin is used as a loading control. Statistical significance was assessed using two-tailed Student’s t-tests after combining data points from all the KL and K cells. ****p<0.0001. Cell viability assay and Western blots are repeated at least twice. FACS analysis is performed once.

**Supplementary Data Fig. 7.**
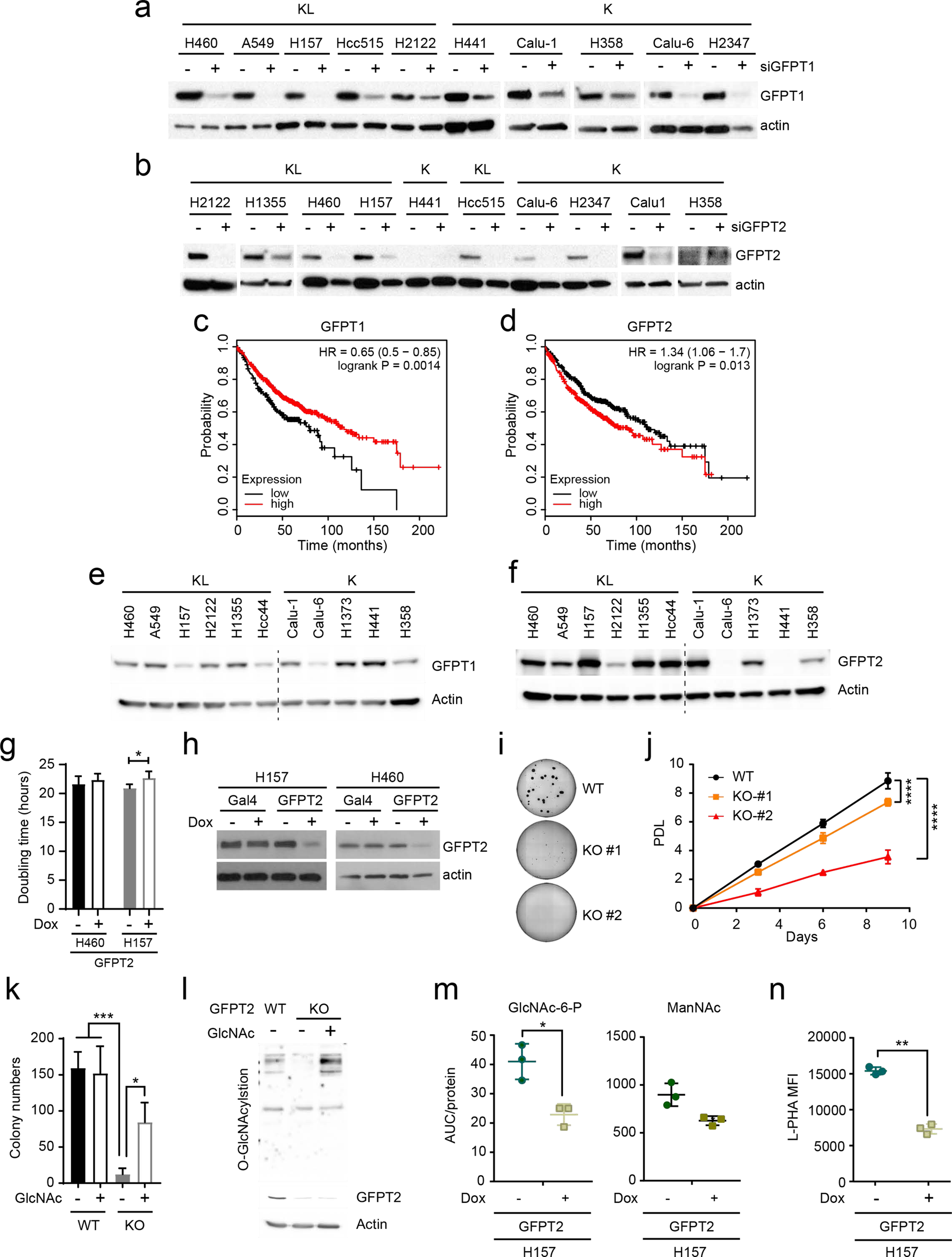
KL cells are sensitive to inhibition of GFPT2. **a,** Abundance of GFPT1 protein in cell lines transfected with a control esiRNA or esiRNA directed against GFPT1. Actin is used as a loading control. **b,** Abundance of GFPT2 protein in cell lines transfected with a control esiRNA or esiRNA directed against GFPT2. Actin is used as a loading control. **c,** Kaplan-Meier plot associating *GFPT1* mRNA expression with survival. Dataset is from KM Plotter (http://kmplot.com/analysis/index.php?p=service&cancer=lung). **d,** Kaplan-Meier plot associating *GFPT2* mRNA expression with survival. **e,** Abundance of GFPT1 in a panel of K and KL cells. Actin is used as a loading control. **f,** Abundance of GFPT2 in a panel of K and KL cells. Actin is used as a loading control. **g,** Effect of Dox-induced GFPT2 deletion on growth in a monolayer culture. Data are the average and SD of 6 replicates. **h,** Abundance of GFPT2 in Dox-inducible GFPT2 KO H157 (left) and H460 (right) cells with or without Dox induction. Actin is used as a loading control. **i,** Representative images of colonies grown in soft agar in Fig. **5g. j**, Effects of a GFPT2 KO on cell proliferation (H460 cells, n = 5). **k,** Effect of GlcNAc on anchorage-independent growth of GFPT2 KO cells. **l,** Global O-GlcNAcylation of GFPT2 WT, KO and KO treated with GlcNAc. **m,** Abundance of GlcNAc-6-P and ManNAc in Dox-inducible GFPT2 KO H157 cells (n=3). **n,** Effect of Dox-inducible GFPT2 KO H157 cells on cell surface L-PHA lectin binding. Statistical significance in **g, j, m,** and **n** was assessed using two-tailed Student’s t-tests. In **j,** to calculate significance on repeated measurements over time, a two-way ANOVA with Tukey’s post hoc test was used. In **k,** statistical significance was assessed using one-way ANOVA with Tukey post hoc test. *p<0.05, **p<0.01, ***p<0.005. Targeted metabolomics was performed once. Soft agar assay, monolayer cell growth, GFPT1 and 2, and O-GlcNAcylation western blotting were assayed twice. All experiments were repeated three times or more.

**Supplementary Data Fig. 8.**
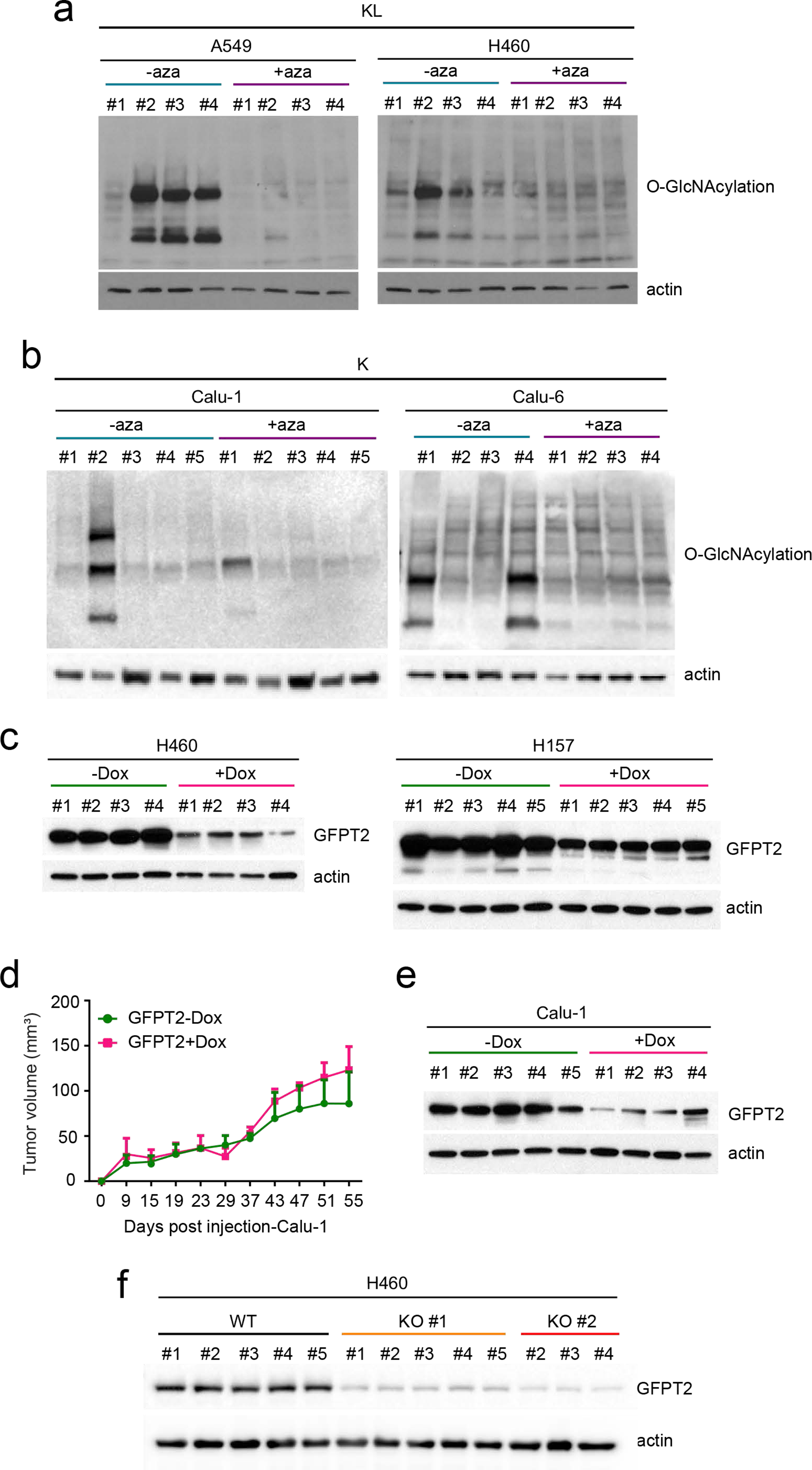
GFPT2 suppression inhibits KL tumor growth. **a,** Global O-GlcNAcylation in A549 (left) and H460 (right) xenografts in presence and absence of azaserine. Actin was used as a loading control. **b,** Global O-GlcNAcylation in Calu-1 (left) and Calu-6 (right) xenografts in presence and absence of azaserine. Actin was used as a loading control. **c,** Abundance of GFPT2 protein in Dox-inducible GFPT2 KO H460 (left) and H157 (right) xenografts with or without Dox induction. Actin was used as a loading control. **d,** Growth of Dox-inducible GFPT2 KO Calu-1 xenografts in presence and absence of Dox. Mean tumor volume and SEM are shown for each group (n=5 for GFPT2-Dox, n=4 for GFPT2+Dox). **e,** Abundance of GFPT2 protein in Dox-inducible GFPT2 KO Calu-1 xenografts with or without Dox induction. Actin was used as a loading control. **f,** Abundance of GFPT2 protein in GFPT2 WT and KO (two independent clones) H460 xenografts with or without Dox induction. Note that only three mice bearing KO #2 cells developed tumors. Actin was used as a loading control. Statistical significance was assessed using two-way ANOVA followed by Sidak’s multiple comparisons test. Mouse experiments were performed once. Western blots were repeated twice.

**Supplementary Data Fig. 9.**
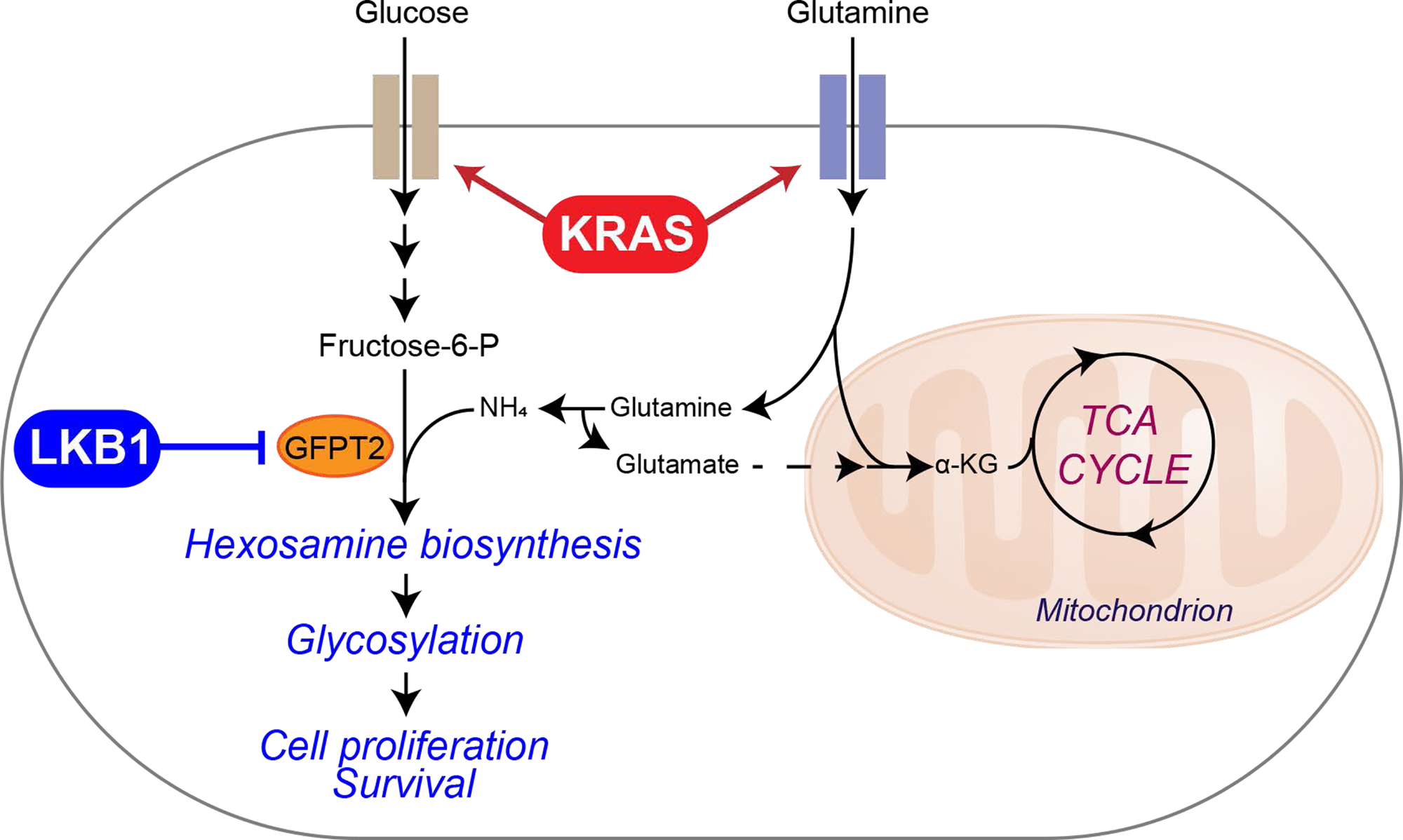
Illustration of the HBP and tricarboxylic acid (TCA) cycle. Metabolic alterations mediated by concurrent mutations of KRAS and LKB1 render cells dependent on GFPT2 for hexosamine synthesis. Uptake of glucose and glutamine is elevated by mutant KRAS, establishing the environment favoring hexosamine synthesis. LKB1 loss in the context of KRAS mutation then enhances HBP through GFPT2. Increased GFPT2 activity may also contribute to the glutamate pool for anaplerosis in the mitochondria. α KG, α-ketoglutaric acid.

**Supplementary Data Table 1. Metabolomics in GEMM tissues.** A set of 305 metabolites was monitored in 10 tumor tissues/genotype and 6 normal lung tissues (see row labeled “Class”).

**Supplementary Information Table 2: Variable Importance in the Projection (VIP) analysis of metabolomic differences between tissues.** Primary data used in the analysis are in Supplementary Information Table 1.

**Supplementary Information Table 3: RNA seq analysis of mouse normal lung and tumor tissues.** mRNA expression of 2602 metabolic genes are shown.

**Supplementary Information Table 4: Gene Set Enrichment Analysis (GSEA) using mouse normal lung and tumor tissues.** Primary data used in the analysis are in Supplementary Information Table 3.

**Supplementary Information Table 5: Leading-edge gene rank of GSEA results.** Leading-edge gene ranks from Supplementary Information Table 4 are shown.

**Supplementary Information Table 6: Information about genetically engineered mouse model (GEMM).** Sex, date of birth (DOB), body weight at the time of treatment are listed.

## Methods

### Cell lines, Culture, and Reagents

All NSCLC cell lines (A549, H1355, Hcc515, H157, H2122, H2030, H460, DFCI-024, H1155, H1373, H2347, H358, H441, H1792, H2291, Calu-1, Calu-6) used in this study were obtained from the Hamon Cancer Center Collection (University of Texas–Southwestern Medical Center). Cells were maintained in RPMI-1640 supplemented with penicillin/streptomycin, and 5% fetal bovine serum (FBS) at 37°C in a humidified atmosphere containing 5% CO_2_ and 95% air. All cell lines have been DNA fingerprinted for provenance using the PowerPlex 1.2 kit (Promega) and were mycoplasma free using the e-Myco kit (Bulldog Bio). H460- and H2122-EV, -LKB1 WT were generated as described previously (CPS1 paper). pBABE-FLAG-LKB1 WT were from Lewis Cantley (Addgene plasmid #8592). GFPT2-deficient H460 clones for population doubling, colony formation, mouse xenograft assays were generated using the original CRISPR-Cas9 systen^43^. To generate inducible CRISPR-Cas9 GFPT2 KO cell lines, parental cells (H460, H157, Calu-1) were first infected by pCW-Cas9 plasmid (Addgene plasmid #50661), sorted by puromycin selection. Stable integrants were then further infected by modified lentiGuide-puro plasmid (Addgene plasmid # 52963) whose original puro selection marker is replaced with ZSGreen1 which is amplified from pLVX-EF1a-IRES-zsGreen1 (Clontech: Catalog No. 631982). lentiGuide-ZSGreen1 expressing sgRNA targeting yeast Gal4 as negative control or sgRNAs targeting GFPT2, and stable integrants were obtained by flow cytometry (FACS Aria II SORP). The primers used to generate sgGFPT2 constructs were as follows:

sgGFPT2-#1 forward, 5-CACCGTACAGAGGCTACGACTCGGC-3,
sgGFPT2-#2 forward, 5-CACCGACTACAGAGTCCCCCGGACG-3,
sgGFPT2-#1 forward, 5-CACCGTCGGCATTGCCCACACGCGC-3,
sgGFPT2-#1 forward, 5-CACCGAGACAGCATGGACTTAAAAG-3.

Doxycycline was from Research Products International (RPI), N-Acetyl glucosamine (GlcNAc), azaserine, OSMI-1, and cisplatin were from Sigma-Aldrich.

### Metabolomics

Mouse tissue metabolomics was performed by Metabolon, Inc. Data from human tissues were retrieved from a previous study^14^. NSCLC cell lines were plated at 3-5 × 10^6^ cells per 10 cm plate for 16hr prior to harvest. Two hours before harvest, cells were incubated with fresh media. At the time of harvest, cells were washed with ice-cold saline, lysed with 80% methanol in water and quickly scraped into an Eppendorf tube followed by three freeze–thaw cycles in liquid nitrogen. The insoluble material was pelleted in a cooled centrifuge (4 °C) and the supernatant was transferred to a new tube and evaporated to dryness using a SpeedVac concentrator (Thermo Savant). Metabolites were reconstituted in 100 µl of 0.03% formic acid in LCMS-grade water, vortex-mixed, and centrifuged to remove debris. For human NSCLC metabolomics, frozen tissues were weighed and divided into 3-9 fragments (∼3mg/each fragment) for technical replicates. Fragments were homogenized in 80% methanol in water and centrifuged at 14,000g for 15 minutes (4 °C). The supernatant was transferred to a new tube and evaporated to dryness as described above for cell lysates. Samples were randomized and blinded prior to analyzing by LC/MS/MS. LC/MS/MS and data acquisition were performed using an AB QTRAP 5500 liquid chromatograph/triple quadrupole mass spectrometer (Applied Biosystems SCIEX, Foster City, CA) as described previously^44^ with injection volume of 20 µL. Chromatogram review and peak area integration were performed using MultiQuant software version 2.1 (Applied Biosystems SCIEX, Foster City, CA), and the peak area for each detected metabolite was normalized against the total ion count (TIC) of that sample to correct any variations introduced from sample handling through instrument analysis. The normalized areas were used as variables for the multivariate and univariate statistical data analysis. All multivariate analyses and modeling on the normalized data were carried out using Metaboanalyst 4.0 (http://www.metaboanalyst.ca). Univariate statistical differences of the metabolites between two groups were analyzed using two-tailed Student’s *t*-test.

### ^15^N glutamine labeling

Cells were plated at 3 × 10^6^ cells per 10cm plate for 16hr prior to labeling. The next day, cells were incubated in labeling media containing 2 mM [γ-^15^N]glutamine for 6hr before harvest. At the time of harvest, the cells were washed with ice-cold saline, lysed with 80% methanol: 20% water and processed as described above (Metabolomics). Metabolites were reconstituted in 100 µl of 0.1% formic acid in LCMS-grade water, vortex-mixed, and centrifuged to remove debris. Samples were randomized and blinded prior to analyzing by LC/MS/MS. LC/MS/MS was performed on an AB SCIEX 5500 QTRAP liquid chromatography/mass spectrometer (Applied Biosystems SCIEX), equipped with a vacuum degasser, a quaternary pump, an autosampler, a thermostatted column compartment, and a triple quadrupole/ion-trap mass spectrometer with electrospray ionization interface, controlled by AB SCIEX Analyst 1.6.1 Software. SeQuant^®^ ZIC^®^-pHILIC (150mm×2.1mm, 5µm) PEEK coated column was used. Solvents for the mobile phase were 10mM NH4Ac in H2O (pH = 9.8, adjusted with conc. NH4OH) (A) and acetonitrile (B). The gradient elution was: 0–15 min, linear gradient 90–30% B; 15–18 min, 30% B, 18–19 min, linear gradient 30–90% B; and finally reconditioning it for 9 min using 90% B. The flow-rate was 0.25 ml/min, and injection volume was 20 µL. Columns were operated at 35°C. Declustering potential (DP), collision energy (CE) and Collision Cell Exit Potential (CXP) were optimized for each metabolite by direct infusion of reference standards using a syringe pump prior to sample analysis. MRM data were acquired with the following transitions: hexosamine-6-phosphate: 258/79 (CE, −40V); N-acetylhexosamine-1/6-phosphate: 300/79 (CE, −80V); Uridine diphosphate N-acetyl hexosamine: 608/204 (DP, 80V; CE, 12V) and 282/79 (DP, −280V, in-source fragmentation; CE, −72V); Neu5NAc: 310/292 (CE, 8V). Chromatogram review and peak area integration were performed using MultiQuant software version 2.1 (AB SCIEX). The peak area for each metabolite was normalized against total cell number. The normalized area values were for statistical analysis.

### Mouse Xenografts and Infusions

Animal procedures were performed with the approval of the UT Southwestern IACUC. Doxycyline-inducible GFPT2 KO cells (H460, H157, and Calu-1) were suspended in RPMI, mixed with Matrigel (Becton Dickinson), and 0.1×10^6^ for H460 and 3×10^6^ cells for H157 and Calu-1 were implanted subcutaneously into 6-week-old NCRNU mice. After tumor cell injection, mice were randomized and then allocated into cages. Mice were fed regular chow or Doxycycline-containing chow (200mg/kg, Bio-Serv), starting 1 day after implantation. For azaserine treatment, cells (H460, A549, Calu-1, or Calu-6) were suspended in serum free RPMI, mixed 1:1 with Matrigel (Becton Dickinson), then 0.1×10^6^ (H460), 1×10^6^ (Calu-6), 2×10^6^ (A549), 3×10^6^ (Calu-1) cells were subcutaneously injected into the flanks of 6-week-old NCRNU mice. After tumor cell injection, mice were randomized, allocated into cages, and intraperitoneally injected azaserine at 2.5mg/kg or PBS when the xenografted tumors measured ∼100mm^3^. Injections were performed every other day for a total of 6 or 7 doses. Tumor size was measured every other day with electronic calipers. Tumor volumes were calculated every 3 to 4 days by caliper measurements of the short (a) and long (b) tumor diameters (volume = a^2^×b/2) or of tumor height (h), short (a) and long (b) tumor width (volume=h×a×b/2) depending on tumor shape. Infusions occurred when flank tumors were 0.5-1.5cm diameter. Mice were fasted the night before the infusion for about 16 hours, and the next day, 25-gauge catheters were placed in the lateral tail vein under anesthesia. [γ ^15^N]glutamine infusions started immediately after implantation of the catheter and continued for 5 hours at the rate of 5mg/kg/min with the total dose of 1.5g/kg in 750µl saline, also under anesthesia. The glutamine solution was administered as a bolus 150µl/min (1min) followed by a continuous rate of 2.5ml/min for 5 hours. Blood samples (10-20µl) were collected at 0.5, 1, 2 and 5 hours via retro-orbital bleed. Animals were euthanized at the end of the infusion, tumors were harvested, rinsed briefly in cold saline and frozen in liquid nitrogen. Frozen tissues were then weighed and divided into 3 fragments for technical replicates. Fragments were homogenized in 80% methanol:20% water and processed and analyzed as described above (Metabolomics).

### Cell growth, Cell death, and Viability

To monitor proliferation in monolayer culture, 1-3 × 10^5^ cells were seeded in a 6 cm dish. Every 3 days, cells were trypsinized and counted with a hemacytometer. The live cell content was estimated using CellTiter-Glo assay (CTG, Promega). To examine cell death, cells were treated as indicated in the Figure Legend and stained with propodium iodide (PI) or with Annexin V-FITC and PI. Cells were then analyzed by flow cytometry (FACS Aria II SORP).

### Soft-Agar Colony Formation Assay

Four days after Dox induction, cells (2,000/well) were suspended in 0.375% agar (Noble agar, Difco) pre-equilibrated with growth medium, over a 0.75% bottom agar layer in each well of a 6-well plate. There was no pre-doxycycline treatment for GFPT2 KO and WT H460 clones generated by the original CRISPR-Cas9 system. Colonies were allowed to form for 20-22 days with intermittent medium supplementation (a few drops twice a week). Images were acquired with G box-Syngene (Syngene) and colonies were detected with GeneTools software (Syngene). To examine constitutive CRISPR-mediated GFPT2 KO clones in soft agar, the same procedures was used.

### Analysis of cell invasion

Invasion assays were carried out by seeding 5 × 10^4^ cells on Corning BioCoat Matrigel Invasion Systems (354480, Corning). The Corning BioCoat control inserts were used as a cell migration control (354578, Corning). Fetal bovine serum (5%) in RPMI 1640 was used as the chemoattractant. Non-invasive cells on the upper membrane of the insert were removed with Q-tips, and invasive cells attached to the bottom membrane of the insert were stained with Hema 3(tm) Fixative and Solutions (Fisher Scientific) after 24-hour incubation at 37°C. Invaded cells were imaged with an Olympus IX81 microscope and quantified using ImageJ.

### FACS analysis

Lectin binding was evaluated by flow cytometry as described previously^45^. The following reagents were used at the final concentrations indicated: allophycocyanin-streptavidin (APC-Strep) (Life Sciences, S-868), 5 ug/mL), LEA-biotin (Vector labs B-1175, 1 µg/mL), L-PHA rhodamine (Vector labs RL-1112, 20 µg/mL). Empty vector (EV) control and LKB1 WT expressing H460 and H2122 cells, and doxycyline-inducible GFPT2 KO H460 and H157 cells were harvested by centrifugation, washed twice with DPBS, then resuspended in DPBS at 2.0 × 10^6^ cells/ml. Then, 200 µL of cell suspension was transferred to V-bottom 96-well plate and pelleted by centrifugation at 600g for 5 minutes. The cell pellets were incubated with 100 µL of lectin diluted in DPBS for 60 min at 4 °C (LEA incubation was 30 min), then washed with DPBS three times. When a secondary detection reagent was used, cells were incubated with 5 µg/mL APC-Strep in DPBS for 45 min at 4 °C. Fluorescence was analyzed by flow cytometry on a FACS Aria II SORP with dual lasers at 488 nm and 635 nm. Plots show the mean fluorescence intensity (MFI), typically for 10,000 cells.

### RNAi

Transient gene-silencing experiments were performed with endoribonuclease-prepared siRNAs (esiRNA, Sigma) for GFPT1 and GFPT2, and with ON-TARGETplus-SMART pools (Dharmacon) for LKB1 and OGT. Viability assays were performed after 96hr and cell death analyses and all other western blots were performed after 48hr.

### Western Blot Analysis

Protein lysates were prepared in either RIPA or CHAPS buffer and quantified using the BCA Protein Assay (Thermo Scientific). Protein was separated on 4%–20% SDS-PAGE gels, transferred to PVDF membranes, and probed with antibodies against O-GlcNAcylation (Santa Cruz, sc-59623), β-actin (Abcam, ab8227), cyclophilin B (clone EPR12703(B), ab178397), CPS1 (ab3682), GFPT1 (ab125069), GFPT2 (ab190966), phospho-ACC (Cell Signaling, #11818), LKB1 (Cell Signaling, #3050), and OGT (Sigma, O6264-200UL).

### Immunohistochemistry

Paraffin-embedded sections from mouse lung tumors were deparaffinized with xylene followed by ethanol rehydration, fixed in 4% paraformaldehyde, and antigen-retrieved with 10 mM sodium citrate pH 6.0. Sections were then subjected to endogenous peroxidase blocking with 0.3% H2O2. Bovine serum albumin (BSA, 3%) in 0.1% PBST was used as the blocking agent and antibody dilution solution. After 1hr blocking, samples were incubated overnight at 4°C with the primary antibody (Galectin-3, ab2785), washed, incubated for 30 minutes with diluted biotinylated secondary antibodies (Vector lab, PK-4000). After washing, sections were incubated for 30 minutes with VECTASTAIN^®^ ABC Reagent, washed for 5 minutes in buffer, incubated in peroxidase substrate solution until desired stain intensity developed. Images were acquired with an Olympus IX81 microscope.

### Patient Survival Data

Differences in survival based on either *GFPT1* or *GFPT2* mRNA expression was determined in lung adenocarcinoma tumors from The Cancer Genome Atlas (TCGA)^46^ and data from KM Plotter (http://kmplot.com/analysis/index.php?p=service&cancer=lung)^47^. Methods for data generation, normalization, and bioinformatics analyses were previously described in the publication. For the present analysis, data from this cohort was analyzed using Lung Cancer Explorer (http://lce.biohpc.swmed.edu).

### Statistical Analysis

No statistical methods were used to predetermine sample size. Metabolomics and flux analysis samples were randomized prior to LC-MS/MS analysis. For xenograft experiments, mice injected with tumor cells were randomized prior to being allocated to cages. All other experiments were not randomized, and the investigators were not blinded to allocation during experiments or to outcome assessment. Experiments in Figs 1a, 1c, 1d, 2b, 2c, 6b, 6c, 6e-f, Supplementary Data Figs 1c, 3a, 5e, 6b, 7m, 8d were performed once. Experiments in Figs 2e, 3b, 3c, 4c, 4e, 5a-i, 6a, 6d, Supplementary Data Figs. 3b, 3c, 6a, 6c, 7a, 7b, 7g, 7j, 7l, 7n, 8a-c, 8e, 8f were performed twice. All other experiments were performed three times or more. Variation for xenograft tumor volume is indicated using standard error of the mean, and variation in all other experiments is indicated using standard deviation. To assess the significance of differences between two conditions, a two-tailed Welch’s unequal variances t-test was used. Where the data points showed skewed distribution (e.g. Supplementary Data Fig. 1c), Wilcoxon signed rank test was performed. For comparisons among three or more groups, a one-way ANOVA followed by Tukey’s multiple comparisons test was performed. To assay significance of differences between K and KL cells on azaserine treatment, R package drc was used to fit a four-parameter log-logistic model for data from each cell line. Area under the fitted curve was calculated for each cell line. Mann-Whitney U test was used to calculate p-value from comparing AUC values from KL cell lines and K cell lines. To examine significance in xenograft tumor growth between two or among three or more groups, two-way ANOVA followed by Tukey’s multiple comparisons test was performed. Before applying ANOVA, we first tested whether there was homogeneity of variation among the groups (as required for ANOVA) using the Brown–Forsythe test. In all xenograft assays, we injected 6-7 week old NCRNU mice (both male and female) 10 mice per treatment, as we expected based on previous pilot experiments to observe differences in tumor size after 2 weeks.

